# Universal genomic constraints in the evolvability of thermal physiology

**DOI:** 10.64898/2025.12.12.693958

**Authors:** Brian P. Waldron, Eliza Tarimo, Palash Sashittal, Noel Naughton, Martha M. Muñoz, Josef C. Uyeda

## Abstract

Thermal physiological traits such as body temperature often show surprisingly slow evolutionary rates over macroevolutionary time, despite apparent lability at microevolutionary time scales. While long-term stabilizing selection may slow rates of thermal evolution, we propose an alternative hypothesis from a bottom-up, population genomic perspective: the nature of body temperature (*T_b_*) as an organism-level trait that must accommodate diverse protein thermal performances leads to evolutionary constraints at the organismal level. We support this hypothesis using a simulation framework in which we modeled and compared the rates of evolution for *T_b_* alongside one or more proteins. Protein performances and organismal *T_b_* were modeled as evolving, QTL-encoded traits, and organismal fitness was determined based on *T_b_* given the performance curves of each protein. As predicted, a greater number of proteins led to drastic decrease in the rate of *T_b_* evolution. When a shift in environmental temperature was simulated, *T_b_* evolved with an initial rapid pulse toward the new optimum, followed by a phase of gradual evolution as the cumulative fitness costs of mismatching *T_b_* and protein optima constrained thermal adaptation. That is, lability and stasis are predictable features of body temperature evolution: rapid, yet bounded microevolutionary bursts followed by long phases of sluggish evolution are both expected outcomes of directional selection operating on hierarchically structured traits like *T_b_*. We suggest that protein thermal coordination might contribute to intrinsic, universal macroevolutionary patterns of stasis in organismal physiology across endotherms and ectotherms.

## Introduction

Through time and across species, phenotypic traits differ greatly in their degree of variation and rates of change. While some traits may be highly labile, others can show exceptionally slow rates of evolution over epochal time scales (1–3). Such cases of slow phenotypic evolution at macroevolutionary scales are often surprising given the rapid evolution observed in many natural and artificial selection experiments, some spanning only a single generation (4–6). This apparent conflict forms the conundrum known as the “paradox of stasis” (7–9). If natural selection has the raw materials—heritable phenotypic variation—to change phenotypes within populations at short time scales, then why do we not observe more trait disparity across the tree of life?

Several ideas have been proposed to bridge the presumed gap between microevolutionary and macroevolutionary patterns and explain the paradox of stasis. These hypotheses include practical issues such as limitations in our ability to measure stabilizing selection (10), but also biologically relevant processes such as developmental constraints (11), long-term maintenance of adaptive zones (12–14), and slow molecular evolution in phenotypically conserved clades (15, 16). Another possibility is that trait evolvability—the capacity to generate heritable variation available for selection (17)—can be constrained when a trait is an emergent phenotype resulting from complex, hierarchical networks of underlying proteins and tissue systems. At a molecular level, epistatic interactions grow increasingly relevant, with sign epistasis (the change in the direction of a mutation’s effect on fitness depending on the state of another locus) causing rugged fitness landscapes that reduce evolvability (18). At an organismal scale, traits that are a part of an emergent phenotype (*e.g*., the shoulder girdle or limbs within the highly modified body plan of turtles) do not evolve in isolation, but must instead evolve together as a series of correlated steps at each component trait, a process called “correlated progression” (19, 20). This obligate coordination results in the reduced evolvability of the system as a whole.

The evolution of thermal physiology presents a clear example of this paradox of enigmatic stasis (21). When measured at short timescales, thermal physiological traits (*e.g*., field body temperature, or critical thermal limits such as *CT_min_*) can appear highly labile. For example, both plastic and adaptive responses of ectotherms, such as squamates (snakes and lizards), can result in rapid shifts in field body temperature (hereafter, *T_b_*) or critical thermal limits of one or several degrees Celsius to environmental variation or climate extremes (22–24); but see (25–27). These short-term responses, however, contrast starkly with the high phylogenetic conservatism across species, with field *T_b_* variation among squamate lineages spanning 14.8 to 40.8°C (28). While this spans a wide range of environmental variation, available thermal environments vary substantially more. For example, across the globe, mean minimum and maximum daily temperatures of the coldest and warmest months, respectively, range from −65.9°C to 47.0°C (29). Given approximately 202 million years (Ma) of evolution, it is perhaps surprising that cumulative microevolution has not resulted in a greater range of *T_b_* across squamates. Notably, macroevolutionary stasis is present to an even greater extent in endotherms such as mammals and birds, in which field *T_b_* among species range approximately 10°C within each group despite ∼218 and ∼107 Ma of divergence, respectively (28). This result is again surprising considering that populations show heritable variation in *T_b_*, and artificial selection experiments in laboratory rats have demonstrated the capacity for rapid directional selection of approximately 0.21–0.23°C in only four generations (30). Adaptation and plasticity can rapidly achieve a substantial fraction of the variation observed across species, so deciphering the macroevolutionary trends requires deeper consideration and new insight.

Similarly slow macroevolutionary rates of *T_b_* evolution between ectotherms and endotherms suggest universal constraints underlying thermal physiological traits. Much previous work has focused on how certain traits, such as upper and lower thermal limits, may have specific fundamental constraints. For example, orthologous proteins from cold-adapted species tend to show greater thermal sensitivity compared to those of warm-adapted species, yet both high and low temperature extremes generally decrease the availability of protein conformations compatible with ligand binding (31, 32). In addition, while lower temperatures may reduce enzymatic activity or the speed of muscular activity, higher temperatures may have a sharper and less evolvable thermal limit at which protein denaturation begins and locomotor function ceases (23, 33, 34).

However, other constraints are possible. One intrinsic constraint is that despite such physiologically different strategies across taxa, all organisms represent hierarchical systems regarding thermal physiology. Most organisms are composed of thousands of proteins that vary in their thermal performance properties—with approximately 20,000 protein-coding loci in humans possibly coding for millions of proteoforms (35)—but typically an organism has effectively only a single body temperature at any given time that all proteins experience (36, 37). Therefore, the evolution of *T_b_* must accommodate temperatures required across the proteome, an intrinsic source of selection that we refer to as protein thermal coordination (Fig. 1). The necessary coordination of *T_b_* and performances among proteins should represent a correlated progression that results in slowed *T_b_* evolution overall (38). As some macroevolutionary studies have found evidence for stasis in thermal traits (28, 33), while others suggest more rapid local adaptation with the phylogenetic effect resulting from niche tracking (38), we sought to test the consequences of the correlated progression model to rates of *T_b_* evolution.

**Figure 1.**
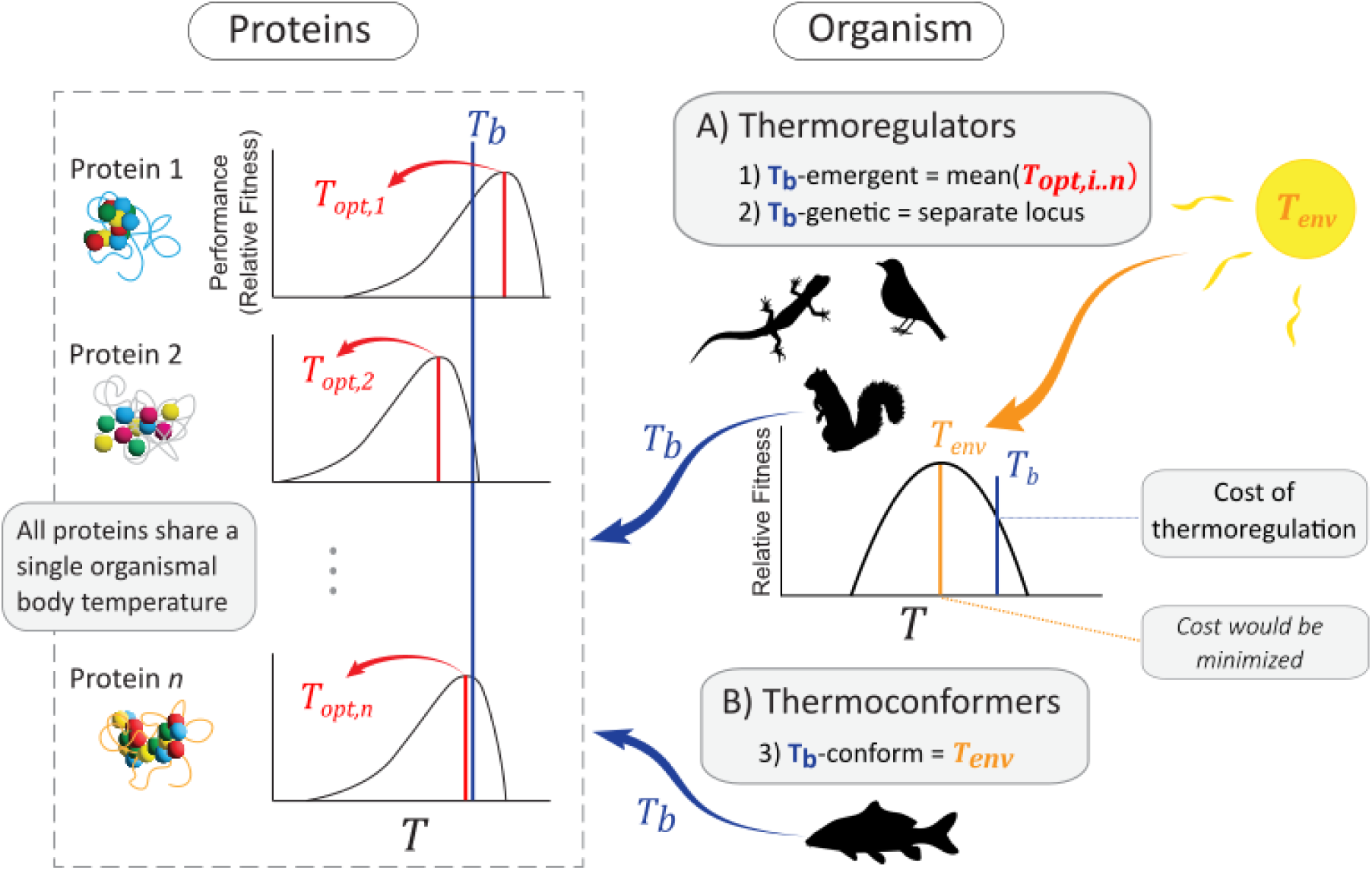
Conceptual model for simulations of body temperature (*T_b_*) evolution. Any organism has a single *T_b_* that all proteins experience (*left*). The contribution of each protein to fitness is determined by a fitness function (*i.e.* thermal performance curve) given *T_b_*. At the organismal level, *T_b_* is determined in several ways (*right*): A) Thermoregulators have a *T_b_* that is effectively independent of environmental temperature due to physiological or behavioral thermoregulation. *T_b_* as a phenotype is either an emergent property—the mean of the individual’s protein thermal optima (*T_opt,i..n_*; “T_b_-emergent” model)—or an independent locus or loci coding for *T_b_* (“T_b_-genetic” model). For either model, a “cost of thermoregulation” can result from organismal *T_b_* mismatching a fixed environmental temperature (*T_env_*). A greater mismatch implies greater cost (*e.g*. metabolic expense of heat production or risk of predation during basking). B) Thermoconformers have a *T_b_* that matches *T_env_* (“T_b_-conform”). Thus, only protein *T_opt_* and the emergent average of these values evolve through time. For all scenarios, organismal relative fitness given *T_b_* is product of relative fitness values across proteins and, for thermoregulators, the organismal cost of thermoregulation. Silhouettes obtained from PhyloPic.org.

Here, we propose a fundamental shift in how we bridge the discordant results between microevolutionary and macroevolutionary studies of thermal physiology. We suggest that thermal physiological traits should be understood as components of a hierarchical system, in which all proteins, regardless of their phenotypic effects—directly related to thermal physiology or not—vary in their thermal performances despite being united under a single organismal body temperature. Thus, rapid evolution in organism-level *T_b_* in response to selection may be decoupled from, yet eventually constrained by, the evolution of thermal optima across the proteome. We predict that increasing genomic complexity in the form of higher numbers of proteins enhances evolutionary stasis of organismal *T_b_* at the population level. We focus here on *T_b_*, but our results have similar implications for the evolution of other organismal thermal traits, such as critical thermal limits. While our models necessarily contain many simplifying assumptions regarding the genetic architecture of thermal physiology and evolutionary pressures impacting the proteome, our primary goal is to identify the influence of a set of universal features of thermal evolution: 1) Protein performance is affected by temperature, and evolution of an individual protein’s thermal optimum is largely independent (non-pleiotropic) from that of other proteins. 2) Proteins, to a first approximation, must share thermal environments within an organism. We explore the implications of these features and how they affect our view of thermal adaptation in nature.

## Modeling Framework

We developed a simulation framework to model the correlated progression of protein thermal performances and organismal *T_b_* that capture key biological features of thermal physiology across endo- and ectothermic animals. We first describe our models generally, including the operationalization of *T_b_* and our assumptions regarding the evolution of endotherms and ectotherms. We then describe the models’ implementation in the software SLiM 4.1 (39), in which we simulated populations of individuals with increasing numbers of proteins, testing the effect of genomic complexity on evolutionary rates and on the capacity to respond to an increase in environmental temperature. Finally, we derive analytical solutions for the estimated rates and constraints of *T_b_* evolution.

### The base model

We begin with a simple hypothetical scenario. Imagine an organism composed of a single protein. The performance of the organism results entirely from the performance of this protein, and the organism’s fitness is directly related to the match between organismal body temperature (*T_b_*) and the protein’s thermal optimum (*T_opt_*), the temperature at which performance is maximized. Body temperature may be an internally regulated temperature in the case of an endotherm, an externally regulated temperature for a thermoregulating ectotherm that it tracks through the environment, or the average environmental temperature itself (*T_env_*) for a thermoconforming ectotherm. In all cases, *T_b_* determines the temperature experienced by the protein, and fitness is maximized when the organism’s *T_b_* matches *T_opt_* for the protein. Evolution of *T_b_* is straightforward in this scenario; as the protein’s *T_opt_* evolves through adaptive or neutral evolution, selection favors *T_b_* that matches or closely tracks *T_opt_* (and vice versa).

Now imagine that the organism is composed of two proteins. As before, the evolving thermal optimum of each protein (*T_opt,i..n_*) presents a moving target for *T_b_*; however, assuming both proteins contribute equally to fitness and have identical, symmetrical performance curves, fitness with respect to *T_b_* will be maximized when *T_b_* is an average of the two proteins’ *T_opt_*. Unless *T_opt_* is identical across loci, the evolution of *T_b_* is now pulled in multiple directions, and neither protein is at its *T_opt_* unless *T_opt,1_ =T_opt,2_*. The rate of evolution for *T_b_* is also expected to slow relative to the single-protein scenario because organismal fitness is no longer isolated to a single trait, instead requiring the concerted evolution of multiple traits (*i.e*., protein thermal coordination). This logic extends to any number of proteins.

### Operationalizing T_b_

We have multiple ways of operationalizing *T_b_* in the model, based on different notions of organismal body temperature’s genetic architecture and evolution. *T_b_* could be an emergent trait, which is simply the average of each individual’s protein *T_opt,i..n_* (our “T_b_-emergent” model; Fig. 1). Such a scenario could resemble a thermoregulating organism whose preferred temperature emerges from its protein performances (40, 41). In this case, *T_b_* is equivalent to an optimal value given variation among protein *T_opt,i..n_*, and organismal fitness is the product of the fitness for each protein given *T_b_* following some fitness function (Fig. 1). A second, more biologically plausible way to operationalize *T_b_* is to treat it as a distinct trait with its own genetic underpinnings (*e.g*., QTLs; our “T_b_-genetic” model; Fig. 1). This second parameterization allows *T_b_* to evolve independently of protein *T_opt,i..n_*. Only overall fitness, and not the trait value of *T_b_* itself, is affected by variation among proteins other than loci specifically underlying *T_b_*. These two models differ in whether *T_b_* is conceptualized as an optimal value that emerges from lower-level performance among one or more proteins, or as a phenotype that can be independently subject to mutation, drift, and selection. A key difference can be illustrated in our simplest, single-protein scenario above: if *T_b_* is emergent, then the protein is by definition always at its *T_opt_*, and evolution proceeds by mutation and drift. If *T_b_* has a distinct genetic basis, protein *T_opt_* and organismal *T_b_* coevolve, with fitness calculated from the mismatch. An alternative model consistent with a thermoconforming ectotherm (“T_b_-conform”) involves the *T_b_* resulting from an externally defined environmental temperature (*T_env_*; Fig. 1) and is described below in the context of thermoregulation.

We pose two generalized thermoregulatory strategies (Fig. 1). First, there is an endotherm or thermoregulating ectotherm (*i.e*., thermoregulators). These organisms maintain their *T_b_* at a consistent temperature, either through internal physiological thermoregulation or behavioral habitat selection on a heterogeneous landscape. The thermoregulating ectotherm is assumed to be a perfect thermoregulator that is always successful at maintaining its preferred temperature; thus, the endotherm and thermoregulating ectotherm are functionally equivalent for our models in that selection on proteins results from *T_b_* independent of environmental variation. Selection at the organismal level, however, caused by *T_b_* mismatching the average environmental temperature (*T_env_*), can result from costs of thermoregulation (Fig. 1), such as risks of predation during basking or the energetic expense of internal heat production (42–44). We assume a greater mismatch between *T_b_* and *T_env_* carries greater fitness costs. Second, we have a thermoconforming ectotherm (*i.e*., thermoconformers), for which we use a modification of our “T_b_-emergent” model (*i.e*., “T_b_-conform”). These organisms have a *T_b_* that matches the environmental temperature (*T_env_*), a value set externally, and it is *T_env_* that all proteins experience. Organismal thermal optima still emerge from average protein *T_opt,i..n_*, although organismal *T_b_* is no longer conceptually equivalent to this value. It is thus the evolution of the organismal optimum (*i.e*., mean of protein *T_opt,i..n_*) that we record for the thermoconforming ectotherm, and we do not expect evolutionary constraints resulting from protein thermal coordination.

### Incorporating thermal performance curves (TPC)

Stabilizing selection is commonly modeled as a symmetrical normal distribution for fitness functions. For thermal traits, however, selection can be modeled following the shape of a thermal performance curve (TPC). TPCs are typically left-skewed in ectotherms, in which performance gradually increases with temperature up to an optimal value, followed by a sharp decline as the increase in the rate of biochemical reactions gives way to protein denaturation. TPCs can evolve for each protein (*T_opt,i_*). By using a TPC as a fitness function, a *T_b_* that is higher than protein *T_opt,i_* contributes more negatively to overall fitness than a *T_b_* that is lower than *T_opt,i_* by the same magnitude. Organismal relative fitness is then calculated as the product of relative fitness across all proteins.

To incorporate a TPC as a fitness function, we used equations from Rezende and Bozinovic (45), which describe performance that increases nearly exponentially with temperature as enzymatic rates increase, followed by a threshold temperature when protein denaturation begins to negatively affect performance:

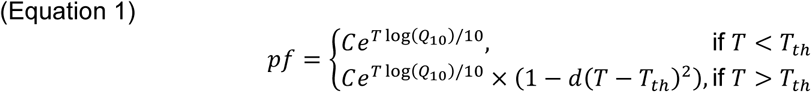

The function has five parameters: *T*, *Q_10_*, *T_th_*, *d*, and *C*. *T* is a given temperature (*i.e*., *T_b_* in our model), and *Q_10_* is the fold change in performance for a change of 10°C in *T*. *T_th_* is the threshold temperature after which a parameter describing the decay rate of performance (*d*) is incorporated into the equation. *C* is a constant that effectively increases or decreases the performance function vertically, although *C* can be ignored here because TPCs were scaled to a maximum of 1.0 when calculating relative fitness. *T_opt_* of the TPC, where performance is maximized, is directly proportional to *T_th_*:

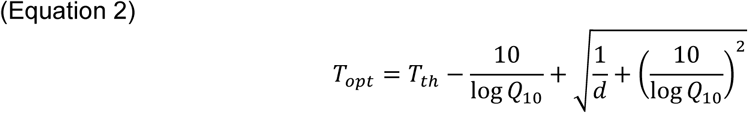

Note that equation 1 implicitly contains a “hotter is better” effect where absolute performance increases as *T_th_* increases (46), but by scaling maximum performance to 1.0, we remove this effect. Our goal here was not to explore the full parameter space of TPCs, but rather to incorporate the general asymmetry of these functions with biologically reasonable parameter values to model relative fitness. For this reason, we allowed only *T_opt_* (and therefore *T_th_*) to evolve, using fixed empirical molecular values of *Q10* (1.813) and *d* (0.0013) from Rezende and Bozinovic (45)(45) for proteins. The consequence of these parameters was small fitness contributions of individual proteins, but potentially large effects across many proteins. For example, the relative fitness for an organism with a *T_b_* one degree higher than *T_opt_* of only a single protein would be 0.996; however, 100 proteins with the same mismatch between *T_opt_* and the *T_b_* would result in a fitness of 0.996^100^ = 0.67.

### Base simulations: no T_env_ or a static T_env_

For our simulations, we used the forward-in-time population genetics simulator, SLiM V. 4.1 (39). All simulations used a Wright-Fisher model of 100,000 generations and a single population composed of *N_e_* = 500 individuals. The genome-wide mutation rate (*μ*) was set at 1 x 10^-7^. We used a *μ* that is substantially faster than empirical values (*e.g*., 0.5 x 10^-9^ for humans) (47), while keeping *N_e_* low to preserve computational performance. To model each protein’s *T_opt_*, we assumed that each protein was underpinned by an independent locus of length 1,000 base pairs, approximating median eukaryotic protein size (353 amino acids) (48). Sites at each locus functioned as quantitative trait loci (QTLs), with mutational effects drawn from a normal distribution of moderate effect sizes N(0, 0.1°C) (49). In all cases, phenotypes were expressed in units of degrees Celsius (°C). To evaluate representative values for the number of proteins, we tested six numbers: 1, 2, 5, 10, 50, and 100. These values are low enough to preserve computational performance, but high enough to allow extrapolation to larger, realistic numbers of proteins. The recombination rate within loci (*i.e*., proteins) was set at *r* = 1 x 10^-8^ for all simulations (50), while different loci were assumed to be independent (*r* = 0.5). Organismal *T_b_* was either calculated as the average of protein phenotypes *T_opt,i..n_* (“T_b_-emergent”), or it had a genotype of either one or five independent loci with the same lengths and effect sizes as our proteins (“T_b_-genetic”). Alternatively, for our thermoconformer model (“T_b_-conform”), we set *T_b_* to equal a fixed environmental value (*T_env_*). Relative fitness was calculated using thermal performance curves as described above.

In our base simulations, the role of environmental temperature varied depending on the thermoregulatory strategy (Fig. 1). For thermoconformers, *T_b_* = *T_env_* was the temperature experienced by all proteins. For thermoregulators, we performed simulations in which *T_env_* was either ignored completely (*i.e*., no *T_env_*, and no organismal costs of thermoregulation), or *T_env_* was included as a static value that the organism experienced, with a greater cost of thermoregulation with greater deviation of *T_b_* from *T_env_*. We used a normally distributed fitness function for the cost of thermoregulation, scaled to a maximum of 1.0, which was multiplied by the relative fitness calculated across proteins. Costs were either “strong” or “weak” (standard deviation of a mean-zero fitness function of 1 or 2), but we note that both values contributed relatively strong effects compared to individual proteins (*i.e*., a 39% or a 12% reduction in relative fitness for a 1°C mismatch of *T_b_* from *T_env_*, respectively).

All simulations began with individuals at the same *T_b_* and protein *T_opt_*: 34.8°C, the optimum for lizard running speed across studies from (45). When *T_env_* was included, it was also maintained at a static value of 34.8°C. Thereafter, evolution proceeded through mutation accumulation, drift, and selection (including stabilizing selection if *T_env_* was incorporated). Mean population *T_b_* was recorded every 1,000 generations, and we estimated variation across replicates using the variance (σ^2^) in final mean population *T_b_* across 100 replicate simulations for each set of parameters. This variance can be considered analogous to Brownian motion evolutionary rates when no *T_env_* was included, as *T_b_* will evolve unbounded with only intrinsic constraints on its rate of evolution. Our estimates can be interpreted as per-year rates based on a one-year generation time but can be easily rescaled by dividing the rate by the generation time.

### Inertia following a shift in environmental temperature

We tested the evolvability of populations in response to an increase in *T_env_*. These simulations began equivalently to each of our static *T_env_* models: we incorporated a 10,000 generation “burn-in” in which *T_env_* remained static at the starting value (34.8°C). Then, we increased *T_env_* by 1°C, where it was held for the remaining 90,000 generations, and directional selection on *T_b_* resulted from costs of thermoregulation. We measured the variation in final mean *T_b_* across replicates (σ^2^), and we also estimated the phylogenetic half-life (t_1/2_) as an estimate of trait inertia. The t_1/2_ measures the average time that a trait takes to evolve from its ancestral state halfway to an optimal value (51); values approaching 0 describe instantaneous adaptation while larger values describe progressively slower adaptation. We estimated t_1/2_ using the following equations (51):

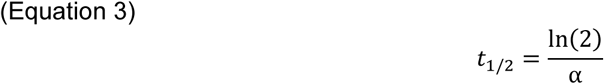

Where α is the strength of the pull towards the phenotypic optimum, which can be defined as:

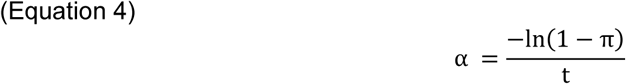

And π is the proportion of the displacement that has been covered at time *t* (i.e. the end of our simulations):

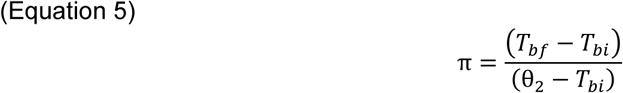

*θ*_2_ represents the shifted environmental temperature, and *T_bf_* and *T_bi_* represent the final and initial body temperatures, respectively. Average values of population *T_bf_* and *T_bi_* across replicates were used in calculations. Here, the ancestral state (*T_bi_*) represented *T_b_* at generation 11,000 soon after the environmental shift. We used generation 11,000 rather than 10,000 when the shift occurred because *T_b_* in our T_b_-genetic models had a rapid initial response to selection followed by more gradual evolution towards the new optimum (see results); therefore, t_1/2_ represents an approximation of the half-life under an OU process for this latter, slow phase after the initial displacement of *T_b_*. We compare our half-lives to macroevolutionary estimates from Tarimo et al. (52), estimated for squamates (76.71 Ma), mammals (63.3 Ma), and birds (49.26 Ma). Mean or median reported generation lengths for each group—4.28, 4.32, and 2.99 years/generation, respectively (53–55)—were used to convert values to units of generations. Finally, we performed simulation sensitivity experiments to assess the effects of modifying mutation rate, effect size, and TPC breadth using both our base model and model with a shift in *T_env_* (details are provided in the SI Appendix, Fig. S1-2).

### Analytical approximations

Finally, we derived analytical solutions for the expected evolution of *T_b_* under our T_b_-genetic and T_b_-emergent models. Namely, we applied a multivariate Ornstein-Uhlenbeck model (56–58) to approximate the rates and constraints of *T_b_* evolution given selection from protein thermal coordination and from the environment through costs of thermoregulation. We detail stochastic differential equations for a multivariate OU model in the SI Appendix. Briefly, we formulated equations that describe the coevolution of *T_b_* and *T_opt_* for each protein. We simplified this multivariate OU process to a bivariate model by tracking mean protein optimum (hereafter, *T_prot_*) instead of each *T_opt_*, assuming a deterministic process in which all proteins respond to selection similarly. To validate the OU model, we performed additional SLiM simulations of the T_b_-genetic model with a 1°C shift in *T_env_* (and “weak” costs of thermoregulation) using a Gaussian equivalent of protein thermal performance curves, allowing us to obtain the linear drift term required for OU models. We calculated additive genetic variance for *T_b_* and *T_prot_* from our simulations and used the width of fitness functions to estimate the strength of selection (*α*; see supplementary appendix 1). Using these values, we estimated changes in *T_b_* and *T_opt_* through time in response to a 1°C shift in the optimum for *T_b_* and compared these to the simulated values.

We also use our simpler T_b_-emergent model to test how a single mutation to a protein in a perfectly coordinated proteome affects evolution as proteome size increases. We derived a series of equations to estimate the mutational “target size,” the effect size for which a mutation is favored by natural selection, under a range of parameters for fitness functions and numbers of proteins (see SI Appendix). We then translate mutational target size to evolutionary rates by deriving approximate evolutionary rates expected under adaptive and drift scenarios. Using biologically plausible parameter values, we then use these formulations test whether decreased mutational target size with higher numbers of proteins can produce evolutionary rates and half-lives consistent with those observed in macroevolutionary studies (52).

## Results

### Base simulations: no T_env_ or a static T_env_

For models lacking any environmental temperature, the final variance in *T_b_* across replicates decreased exponentially with an exponential increase in the number of proteins (Fig. 2A). Here, the variance across replicates can be interpreted as a rate of evolution because selection was affected only by the mismatch between *T_b_* and protein *T_opt_*. These trends were similar regardless of whether *T_b_* was an emergent property (T_b_-emergent) or a trait encoded by one or five loci independent from the proteins (T_b_-genetic). A static environmental temperature, in which thermoregulators suffered additional fitness costs by *T_b_* mismatching the environment, weakened or eliminated the negative relationship between variance and the number of proteins given the strong effects of stabilizing selection from the costs of thermoregulation (Fig. 2B). The thermoconformer model showed a different pattern (T_b_-conform; Fig. 2B); only average protein *T_opt_* was recorded because *T_b_* was equivalent to *T_env_*. Only the relatively weak fitness effects from proteins mismatching *T_env_*, and no costs of thermoregulation, were included for conformers; therefore, the variation across replicates decreased exponentially as the number of proteins exponentially increased (Fig. 2B).

**Figure 2.**
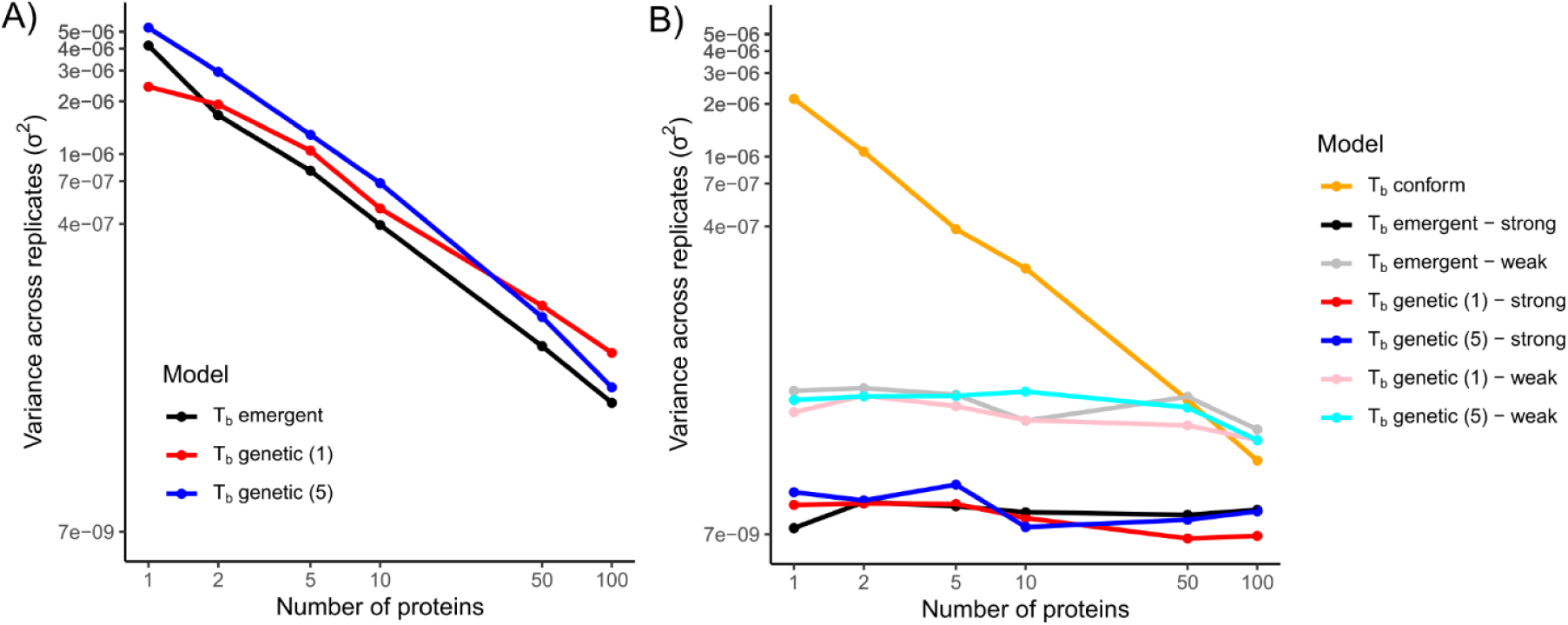
Variance in final population mean body temperature (*T_b_*) across 100 simulated replicates for each model and number of proteins. Models in (A) lack any environmental temperature, and variance across replicates can be conceived as a Brownian Motion rate of evolution (σ^2^) in the absence of external directional or stabilizing selection, whereas models in (B) included a static environmental temperature that introduces stabilizing selection via costs of thermoregulation. Note the log_10_-transformed x- and y-axes shown with original units. Legend values in parentheses for T_b_-genetic models indicate whether one or five loci were used to control organismal *T_b_*. “Strong” and “weak” refer to the cost of thermoregulation, *i.e.* whether fitness functions for organisms mismatching environmental temperature used a mean-zero normal distribution with a standard deviation of one or two. In the T_b_-conform model in (B), variance across final mean protein *T_opt_*, rather than a *T_b_*, was recorded (see text).

### Response to shift in T_env_

Inertia of *T_b_* in response to a 1°C shift in *T_env_* increased as the number of proteins increased for the T_b_-emergent and T_b_-genetic models (Fig. 3A). Rates and half-lives produced by our analytical approximations were consistent with empirical values for birds, mammals, and squamates (Fig. 3B; See *Analytical Approximations* below).

**Figure 3.**
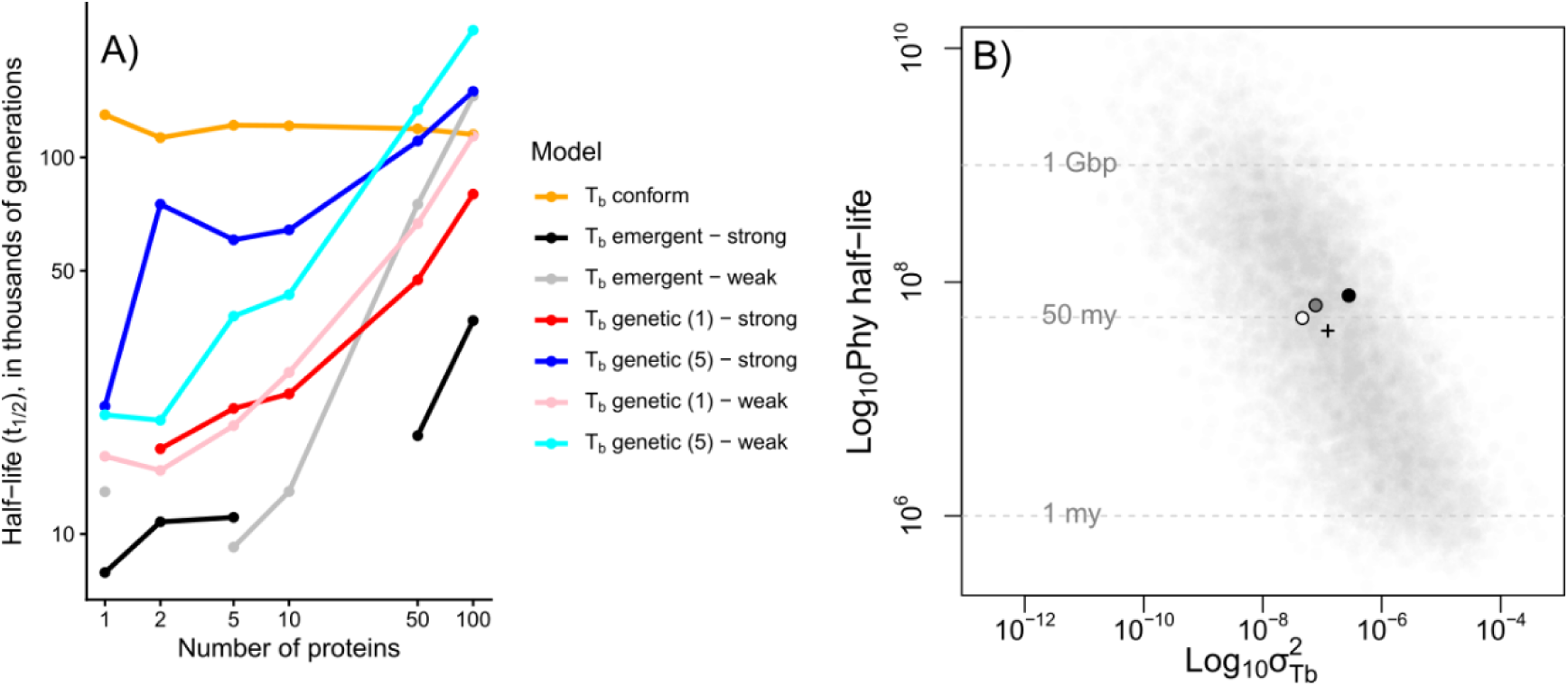
Protein thermal coordination causes evolutionary constraints that increase with greater numbers of proteins and is capable of producing rates and constraints consistent with macroevolutionary studies. (A) Phylogenetic half-life of population mean body temperature (*T_b_*) for SLiM models that included a 1°C increase in environmental temperature partway into the simulation. Note the log_10_-transformed axes shown with original units; the y-axis represents units of thousands of generations. Some models did not have half-lives because the final mean *T_b_* slightly exceeded the new environmental temperature. (B) Macroevolutionary parameters for evolutionary dynamics under an adaptation-inertia model (see *Analytical Approximation*s) estimated from squamates (black), mammals (gray), and birds (white) against 10,000 Monte Carlo simulations (gray dots) over log_10_*N_e_* ∼ logNorm(5, 0.15), log_10_*v ∼ U*(0, 0.036), generation time ∼ logNorm(log(5yrs), 0.25), Number of proteins ∼ *U*(10000, 30000), and *p_nn_* ∼ _U_(0, 0.036).

In the T_b_-genetic model, inertia followed a rapid initial response to selection, in which the costs of thermoregulation strongly pulled *T_b_* towards the new *T_env_* (Fig. 4); however, the benefits of *T_b_* approaching the new *T_env_* were eventually offset by the mismatch between *T_b_* and the proteins’ optima, particularly with greater numbers of proteins (Fig. 4). In other words, because *T_b_* was decoupled from the average of the proteins’ *T_opt_* values, *T_b_* could evolve rapidly until it reached a point when selection due to costs of thermoregulation was as low as the cumulative deleterious effects on protein performances. Slow evolution of *T_b_* and proteins followed thereafter (see also Fig. S3 in the SI Appendix). With our lower value of natural selection from costs of thermoregulation (*i.e*., a 12% reduction in fitness when *T_b_* mismatches *T_env_* by 1°C), mean population *T_b_* never fully evolved to reach a 1°C shift in *T_env_* given >50 proteins (Fig. 4). Instead, we observe rapid evolution in *T_b_* in response to the environmental shift, yet evolution stalls thereafter, on average increasing only 0.56°C at the end of the simulations for 100 proteins. Even with our strongest costs of thermoregulation (*i.e*., a 39% reduction in fitness when *T_b_* mismatches *T_env_* by 1°C), *T_b_* increased only 0.84°C at the end of the simulations for 100 proteins (SI Appendix, Fig. S4).

**Figure 4.**
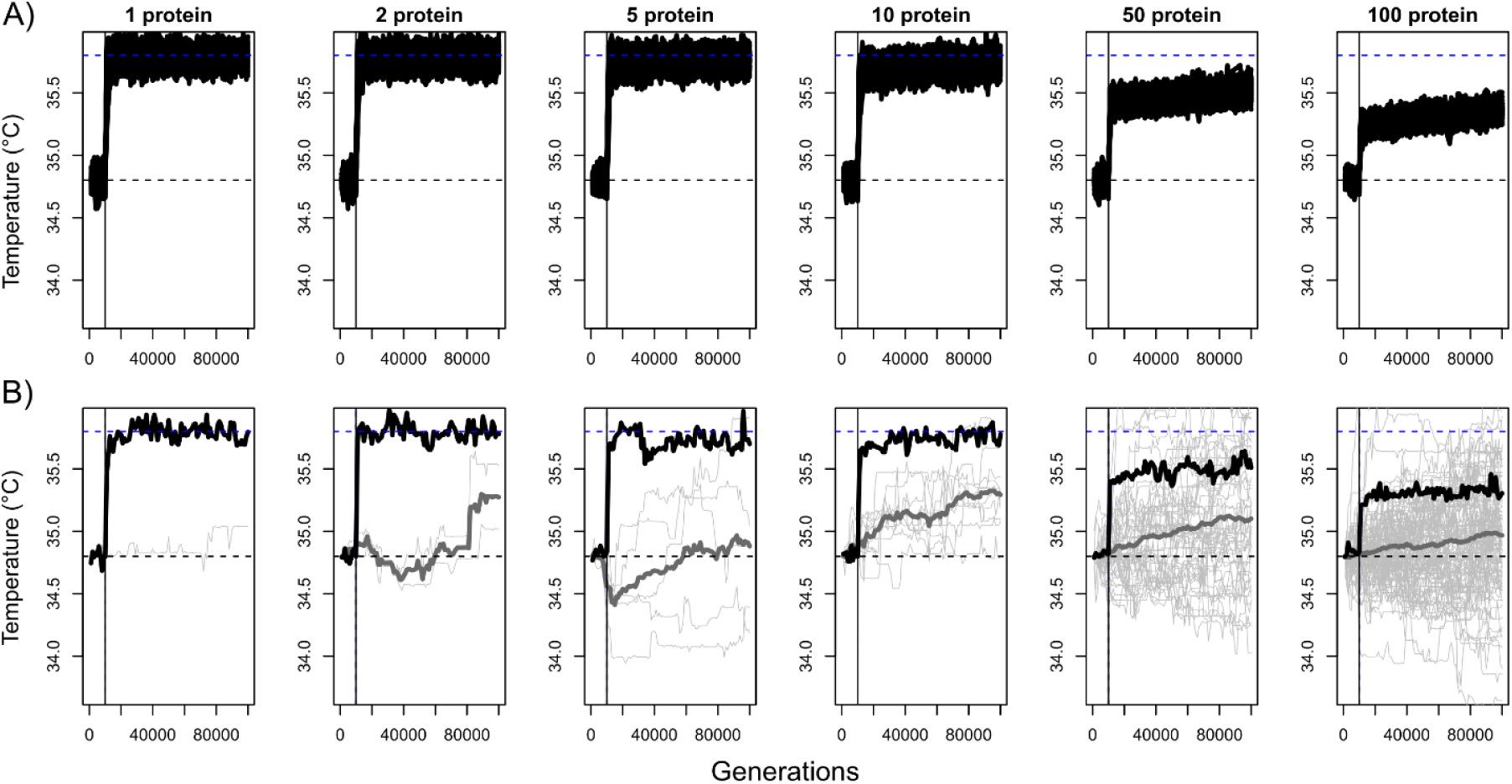
Simulations with *T_b_* coded as a separate genetic trait resulted in two distinct phases of *T_b_* evolution following a 1°C increase in environmental temperature (*T_env_*): a rapid initial evolution in mean *T_b_*, followed by a more gradual increase, particularly with higher numbers of proteins (≥50). Shown here are results for the T_b_-genetic model (five loci for *T_b_*) and our weaker cost of thermoregulation pulling *T_b_* towards the new *T_env_*: (A) Mean *T_b_* across 100 simulated replicates (black lines). (B) An example simulation for each number of proteins showing population mean *T_b_* (black line), mean thermal optima for each protein (*T_opt,i..n_*; thin gray lines), and the mean thermal optima across all proteins (gray line). For all plots, the horizontal dashed gray line represents the starting value of *T_b_*, *T_env_*, and *T_opt,i..n_*, and the horizontal dashed blue line represents the new *T_env_* starting at generation 10,000 (vertical black line).

For the T_b_-emergent model, evolution of *T_b_* lacked the rapid initial response to selection seen in the T_b_-genetic model because of the coupling of *T_b_* with the underlying protein genotypes; selection could not favor an increase in *T_b_* independently from the proteins from which it emerged (SI Appendix, Fig. S5). However, half-lives were comparable to the T_b_-genetic model (Fig. 3A), and *T_b_* failed to ever fully reach the new *T_env_* with 100 proteins (Fig. S5).

Half-lives for the T_b_-conform model, for which only mean *T_opt_* across proteins were recorded, were largely unaffected by numbers of proteins (Fig. 3A). While variance across replicates decreased with more proteins (SI Appendix, Fig. S1), the final mean *T_opt_* across replicates remained consistent (Fig. S6). In the T_b_-conform model, all proteins are effectively under directional selection independently towards the new *T_env_*, with no intrinsic selection against deviating from the mean. Thus, the constraints present when *T_b_* is a trait independent of *T_env_*, as in our other models, are eliminated.

### Analytical approximations

We compared our SLiM simulations including a shift in *T_env_* to predictions derived from a bivariate OU process, in which the strengths of selection (*α*) on *T_b_* and mean thermal optimum across proteins (*T_prot_*) were calculated from simulated additive genetic variance and the widths of fitness functions (see SI Appendix). Predicted values from the deterministic, bivariate OU model provided a good fit to our SLiM simulations, in which both *T_b_* and *T_prot_* through time fell within the range of simulated values (Fig. 5).

**Figure 5.**
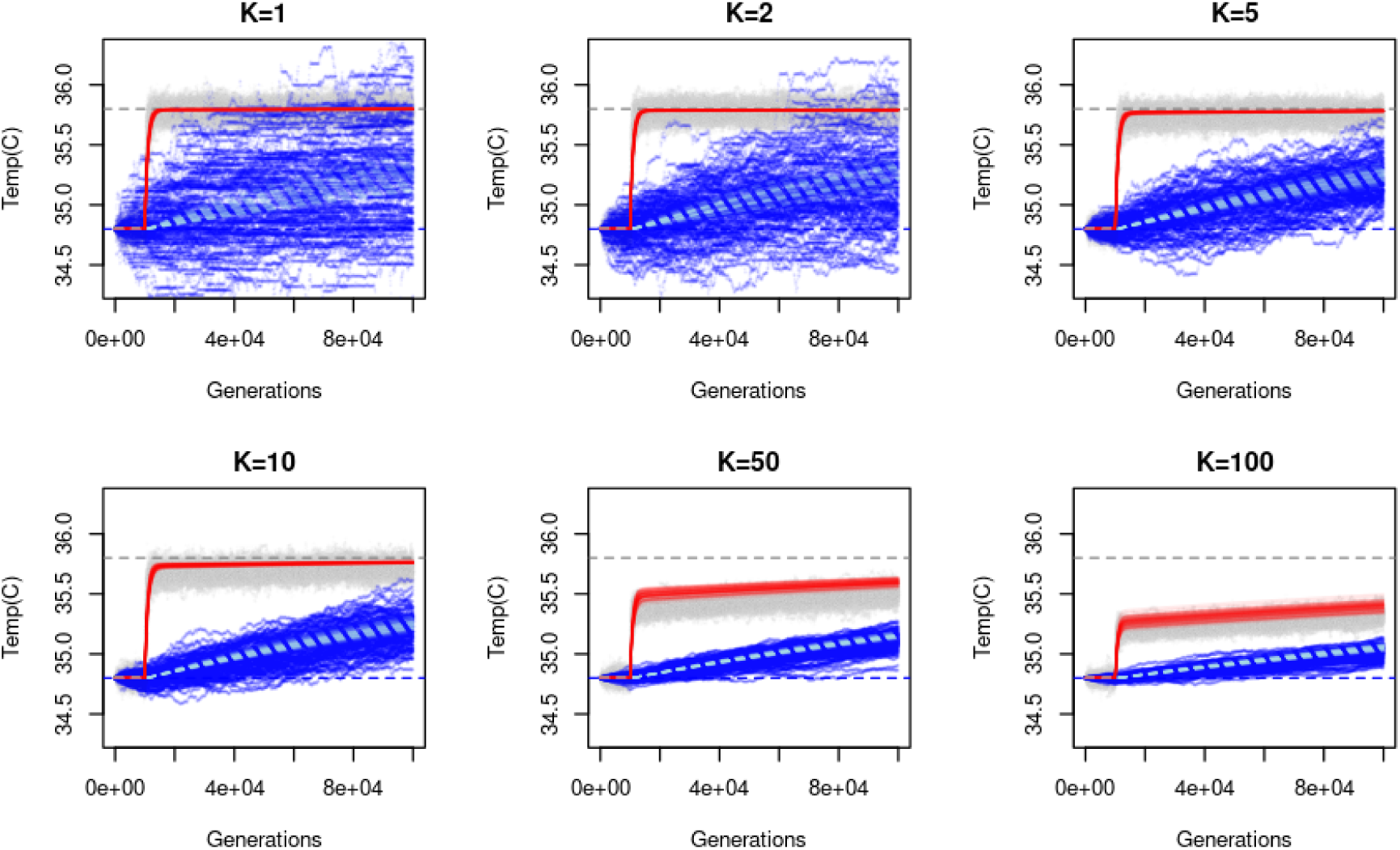
Simulation results for body temperature (gray) and proteome average optimum (dark blue) for different proteome sizes (K) assuming Gaussian thermal performance curves. Solid lines represent expectations from deterministic multivariate Ornstein-Uhlenbeck models parameterized with average genetic variances obtained from individual SLiM simulations for body temperature (red) and proteome optimal temperature (light blue).

Finally, under a T_b_-emergent model, we show that mutational target size drastically decreases for proteomic scales (*e.g.* 10,000 – 30,000 proteins), such that fixation rates approach that of neutral genetic drift (see SI Appendix; Fig. S7). We find that the relationship between mutational target size and proteome size scales inversely with a slope of −1 on the log-log scale (see SI Appendix, Fig. S7). This allows an approximation of the rate of adaptation dependent on the mean of the neutral mutational target window under a model of Brownian motion with a trend. When we simulated evolutionary rates and half-lives under a range of plausible parameter values, including the proportion of nearly-neutral mutations based on experimental values of protein melting temperatures (59), we find that empirical values of evolutionary rates and half-lives from Tarimo et al.(52) are found near the mean of simulated values (Fig. 3B). This suggests that the decrease in mutational target size under protein thermal coordination is capable of producing patterns observed at macroevolutionary scales.

## Discussion

The evolution of thermal physiology exhibits a paradoxical combination of short-term lability and macroevolutionary constraint. In this study, we applied a simulation framework to model the coordinated evolution of proteins and organismal *T_b_*, testing the consequences of protein thermal coordination on rates of short- and long-term evolution. Our models are necessarily simplified, yet they capture a real biological problem: the independent evolution of large numbers of proteins all must operate within the thermal constraints of individual organisms. We found that increasing numbers of proteins decreased the rates of organismal *T_b_* evolution. This resulted from the increasingly multiplicative fitness costs of a single organismal *T_b_* mismatching the *T_opt_* of a greater number of proteins, which continuously acquired mutations that changed their thermal performances. When we incorporated directional selection by adding a shift in environmental temperature, *T_b_* also experienced greater evolutionary stasis as the number of proteins increased, even with relatively strong selection on *T_b_* assuming organismal costs of thermoregulation. Furthermore, a model in which *T_b_* was coded as a separate locus or loci (T_b_-genetic) showed that a rapid initial response of *T_b_* to an environmental shift could quickly reach the limits of protein performance, requiring more gradual correlated evolution of *T_b_* and proteins thereafter. Specifically, the negative fitness effects of reduced thermal protein performance, particularly when cumulative over the thousands of proteins expressed in real organisms, could present significant constraints on the evolvability of organismal thermal physiology. The apparent paradox of thermal lability/stasis is actually an expected product of thermal evolution at different temporal scales: while lability of *T_b_* may be inferred in microevolutionary studies involving relatively small changes in temperature, macroevolutionary stasis is an expected consequence of protein thermal coordination at species’ thermal extremes.

### Implications for rates of evolution: microevolution versus macroevolution

The evolvability of *T_b_* under models with either emergent or genetic origins suggested that adaptive inertia is expected following a shift in environmental temperature. However, the genetic model (*i.e*., T_b_-genetic), in which *T_b_* could evolve independently of the phenotypes of proteins, illustrated how rapid microevolutionary responses of *T_b_* observed in short-term experiments might be compatible with macroevolutionary patterns of stasis. Evolution in response to the shift in *T_env_* included two distinct phases: 1) a rapid initial shift in mean *T_b_* approaching the new *T_env_*, with less absolute change in *T_b_* as the number of proteins increased, followed by 2) a more gradual evolution of *T_b_* towards *T_env_* resembling an Ornstein–Uhlenbeck process (51, 60), with stronger inertia as the number of proteins increased. Put another way, organismal *T_b_*, when genetically decoupled from the *T_opt_* of proteins, could rapidly evolve in response to strong environmental selection pressures, but a lagging evolutionary response of proteins eventually constrains further *T_b_* evolution.

By analogy, imagine attempting to paddle a fishing boat (*T_b_*) towards the shore (*T_env_*) while dozens of fish (*T_opt,i..n_*) are each attached by fishing lines scattered around the boat. The boat may initially paddle easily, but quickly the fish will become aligned on one side of the boat, collectively resisting in the opposite direction. Eventually the force of paddling will cancel out with the cumulative resistance of the fish, and the boat will not move; any more forceful paddling would break a line, losing a fish. Only the sporadic movements of fish in the right direction will ease tension on the lines and allow progressive movement towards the shore.

We speculate that this two-phase process contributes to the apparent paradox of inconsistent rates of physiological evolution across time scales. Natural selection experiments, when focused on evolution of *T_b_* or other organismal traits (*e.g*., *T_pref_*, *CT_max_*, or *CT_min_*), are likely to capture the first phase in which organismal traits may respond rapidly to environmental shifts (24, 61–63). These evolutionary responses may occur with little detrimental effect on much of the proteome if the new *T_b_*, for example, remains within a broad window of competent performance for most proteins; alternatively, the evolution of *T_b_* may occur alongside a major cost in performance across proteins. Organisms are not necessarily optimized for their environments (64), and it is possible that proteins working in suboptimal temperatures is the norm rather than the exception. Nocturnal geckos, for example, are often active in the field at temperatures well below their preferred temperatures measured using thermal gradient experiments (65–67). In the short term, reduced thermal performance of some or even most proteins may be necessary to maintain more important physiological or ecological functions. In other words, rapid microevolutionary adaptation may reflect a balance between protein sub-optimality and ecological-based fitness.

Macroevolutionary studies that report slow rates of physiological or niche evolution, on the other hand, are more likely encounter the second, much longer phase of thermal physiological evolution, in which organismal physiology approaches the limits of protein performance (68–70). In this stage, collective evolution of proteins is necessary to allow organismal *T_b_* to evolve. This process may resemble a correlated progression in which evolution proceeds via incremental changes among proteins because any single protein can only evolve within the constraints of the organismal temperatures, which are themselves constrained by the proteome (19, 20). If a single protein acquires a mutation that shifts its *T_opt_* completely towards a new environmental optimum and *T_b_*, for example, the organism will still suffer a fitness cost with most of its proteome mismatching *T_b_*. Thus, *T_b_* and all proteins are expected to evolve in a stepwise manner once directional selection has reached the limits of performance across the proteome.

While we demonstrate a constraining effect of protein thermal coordination, other factors in addition to the correlated progression of *T_b_* and proteins limit macroevolutionary rates (34). The evolution of upper thermal limits (*e.g*., *CT_max_*) tends to be much slower and more constrained than lower thermal limits (3, 33), suggesting that more specific features of molecular evolution, such as the difficulty of evolving hyperthermostability, set hard limits to evolution (33, 71). In these cases, other macroevolutionary models of physiological evolution may be more appropriate, such as bounded Brownian motion, which can distinguish slow rates of evolution from sharp evolutionary limits (72, 73). A biogeographic alternative is that most vertebrate evolution has occurred in warm climates and regions, and thus the macroevolutionary “constraints” observed in *CT_max_* or *T_b_* may simply reflect environmental legacies (33). These hypotheses are put forth to answer whether we expect more or less divergence in thermal traits between species than observed, given the amount of divergence between species. Our models present an important alternative (but not mutually exclusive) hypothesis that, even in the absence of functional limits to molecular evolution, we might still expect strong evolutionary constraints for organismal thermal physiology.

### Evolutionary stasis in hierarchical traits

Organismal thermal physiology in the form of single, measurable quantities such as *T_b_* represents one example of a hierarchically structured trait. *T_b_* is not only itself the outcome of a variety of traits (*e.g*., metabolism and insulation), but it also directly affects the performance of numerous molecular processes (*e.g*., enzymatic activity). Similar hierarchical systems are common in biology. Body size, for example, which could itself be under directional selection, requires coordinated evolutionary changes in the allometric scaling throughout development, including “biological size”—the relationship between cell size/number and the body size of organisms (74, 75). Body size is also an interesting illustration in how short-term directional selection could be rapid, as observed in the diversity of body sizes (and shapes) of dog breeds following artificial selection, but apparently more constrained across longer time scales (76, 77). Body size, like *T_b_* in our models, may be able to evolve rapidly if selection for the trait is strong, but could ultimately be constrained by negative consequences for the development or performance of many traits at lower levels of organization. However, unlike body size, which may have mechanisms such as pleiotropy or expression-related pathways to coordinate growth, there is no obvious mechanism to change the thermal tolerances of thousands of proteins through a smaller number of mutations.

The evolvability of hierarchical traits is inherently conditioned on the ability to respond to directional selection in the face of stabilizing selection on other trait combinations (78). The efficiency of directional selection for traits such as *T_b_* or body size depends on the degree to which the trait value remains compatible with other traits in the organismal system. Epistatic interactions between mutations also underlie these conditional effects, particularly in the form of sign epistasis, where the sign of its fitness effect—beneficial or deleterious—depends on the genetic background (18, 79). For example, consider a beneficial mutation that increases a protein’s *T_opt_* to a higher value better matched to the environmental optimum, *T_env_* (*e.g*., given *T_opt_* = 29°C and *T_env_* = 30°C, *T_opt_* increases to 29.5°C). Whether this mutation is beneficial or not is conditional upon the genetic background. A proteome where all other proteins have *T_opt_* = 30°C would result in the mutation being universally favored. However, if all other proteins have optima at *T_opt_* = 29°C, the mutation is likely deleterious because any *T_b_* will be mismatched with either the mutant, or the rest of the proteome. Such epistatic interactions grow increasingly complex as the number of independent traits involved in the genetic background increases, further constraining the routes by which populations can navigate an the adaptive landscape that requires tight coordination among these independent components (18).

Note that our findings superficially align with Orr’s (80) “cost of complexity” under Fisher’s geometric model. Like Orr (80), we find adaptation slows with an increasing number of traits that can be modeled via a linear approximation that approaches *K^-1^* for the T_b_-emergent model (Fig. S7). However, for Fisher’s geometric model, mutations are universally pleiotropic whereas here they are on the opposite end of the pleiotropic spectrum—each mutation affects only one protein’s thermal optimum. Furthermore, Orr’s model imposes no absolute constraints on mutational effect sizes and rates of adaptation will scale proportionally with mutational effect size, whereas the hierarchical nature of temperature integration across the proteome makes rates of adaptation largely insensitive to mutational effect sizes (Figs. S1 & S7). Both frameworks, nonetheless, reveal emergent costs that arise from a mismatch between the dimensionality of the evolving system and the low-dimensional vector of adaptation—but in our case, that mismatch combined with hierarchical trait integration imposes an absolute limit on evolvability.

### Thermal physiology across levels of organization

The *T_b_* of the organism that is typically measured by biologists is also the temperature experienced by thousands of proteins throughout the cells, tissues, and organs (exceptions discussed below). Despite the importance of *T_b_* as a trait regularly measured in thermal physiological studies, *T_b_* does not have a simple, well-understood genetic basis (81, 82), reflecting, in part, that body temperature is the outcome of a variety of traits and processes within and between species. Thermoregulating ectotherms require environmental sources of heating and cooling, therefore necessitating behavioral traits, such as preferred temperatures (*T_pref_*), that could have a complex genetic basis with sensory and cognitive components (83, 84).

Thermoregulation can be further aided by morphological characteristics such as color adaptations that maximize heat absorption during basking (85, 86). Endotherms, while also employing behavioral thermoregulation, primarily maintain *T_b_* through the production of metabolic heat and insulation (87, 88). For either case, the optimal organismal *T_b_* must not only maximize the performance of ecological functions, such as foraging or escaping predation, but also facilitate protein performance throughout the body.

We presented two scenarios of how “body temperature” exists as a trait. Our T_b_-emergent model, which treats organismal *T_b_* as the average of protein *T_opt_*, may correspond to the fact that preferred temperatures sometimes have an underlying relationship with protein activity temperatures (40). Our alternative T_b_-genetic model represented a scenario in which organismal *T_b_* is controlled by one or several loci, independent of the genotypes of its remaining proteins (although fitness is dependent on those values). We suggest that the T_b_-genetic model is likely more realistic, and in fact probably underestimates the number of loci involved in producing *T_b_*. Nevertheless, this model may capture how physiological traits that we measure in the lab or field, such as *T_b_*, can evolve independently given selection on specific phenotypes such as metabolic heat production, heat shock proteins, or cognitive components of thermal habitat selection, even if the proteome evolves very little (23).

There are, of course, examples of thermoregulating organisms for which internal temperature varies considerably throughout the body. In addition to surface versus core temperature variation (89, 90), a notable case is of descended testes in mammals, in which the scrotum maintains the testes at a temperature a few degrees cooler than the core temperature (88). Many alternative hypotheses exist for the evolutionary origins of descended testes (91), but species with the trait have reduced sperm output and motility when exposed to higher testicular temperatures (88). This example serves as an exception that proves the rule of organisms effectively having a single *T_b_*: the presence of different body temperatures for different organs must have sufficiently high fitness benefits to outweigh costs of this modularity, such as the developmental risks associated with testicular descent (91).

*T_b_* can also vary temporally, both in daily and seasonal cycles (92, 93). Moreover, *T_b_* often varies throughout life history—female lizards commonly seek higher or lower temperatures during reproduction (94, 95)—or during infection (*e.g*., metabolic fever and behavioral fever) (96). However, while the performances some activities are likely increased or maximized through this temporal variation, other functions are diminished due to fitness trade-offs (97). Therefore, although we modeled fixed body and environmental temperature for each generation, we expect similar constraints on *T_b_* evolution even in the presence of dynamic temperatures.

### Assumptions and extensions of the model

Although the goal of our simulations was not to fit or precisely replicate empirical rates of evolution, which would require more precise empirical estimates of mutational effects and connections between performance and fitness, we sought to describe the general effects of genomic complexity on evolvability and to evaluate its plausibility in explaining macroevolutionary patterns of thermal evolution. Our models incorporated a number of assumptions that will affect the observed rates and constraints of evolution (44). Fitness was assumed to be directly related to thermal performance, multiplicative across the fitness values of each protein, and relative rather than absolute. Using relative fitness with fixed population sizes has the important consequence that extinction is impossible even if the entire population is poorly adapted to a shifting environmental temperature. Applying non-Wright-Fisher models would be a natural extension to account for the possibility of severe population declines in dynamic environments (49). We also used fitness functions that were based upon equally wide thermal performance curves for proteins, or normal distributions for organismal costs of thermoregulation, in each case scaled to a maximum fitness of 1.0. For proteins, we therefore did not consider “hotter is better” effects, in which absolute performance might increase as the thermal optimum increase (46), although as noted above, other factors seem to limit populations from continuously evolving higher *T_b_* and *CT_max_*.

Another important assumption was modeling proteins as independent sequences of linked QTLs that affect their *T_opt_*. We suggest that this is a reasonable model of the way that molecular evolution of protein sequences could have negligible or large quantitative effects on thermal performance, even if the mechanism of thermal evolution involves other factors such as protein chain length (98). However, models including more explicit parameterization of how mutations affect thermal properties given amino acid sequences and folding structure, even for a targeted set of proteins, would help ground the relative trade-offs between the evolution of organismal traits and proteins (99, 100). We implemented relatively weak fitness effects of individual proteins, which here illustrated the point that constraints on *T_b_* evolution could be due to small effects across large numbers of proteins rather than a single protein with large fitness effects, even though large effects are possible (44, 101).

Similarly, a rigorous quantification of the fitness costs of thermoregulation at the organismal level would be a valuable extension of these models. We assumed strong fitness effects with normal distributions (implying that with respect to *T_b_*, it is best to match the environmental temperature), and these values greatly affect the balance between the deleterious consequences of *T_b_* mismatching protein optima versus *T_b_* mismatching environmental temperatures. For most species, the fitness costs of thermoregulation are less clear, as well as how these values relate to protein-level costs. None of these assumptions are likely to change the core result that organismal evolution is constrained to some degree when a single organismal-level phenotype affects the performance of a large number of other traits that vary in their phenotypes. Nevertheless, the expected strength of selection and rates of evolution will depend heavily on the trade-offs between selection at the levels of organisms and proteins.

### Conclusions

Many species have evolved to tolerate thermal extremes. Among vertebrates, some *Liolaemis* lizards from the Andean foothills exhibit a *CT_max_* of over 48°C (102, 103) while the Siberian salamander (*Salamandrella keyserlingi*) can overwinter in freezing temperatures as low as −35°C (104). However, active body temperatures are far more constrained than these tolerance extremes, exhibiting evolutionary stasis and strong inertia that is widespread across taxa. We demonstrated that short-term lability and long-term stasis, to some degree, are expected outcomes of selection acting on a complex, hierarchically structured trait, providing some resolution to the apparent paradox between micro- and macroevolution. This outcome represents a universal constraint that will be common to species with diverse physiological strategies. We envision that these results will provide thermal physiologists with a framework with which to understand compensatory responses to responses to climate change and reconcile these patterns with long-term trends in climatic niche evolution.

## Acknowledgments

We thank the Uyeda and Muñoz labs for comments on an earlier version of the manuscript. We thank the Virginia Tech Center for Mathematical Biology for catalyzing collaboration leading to new mathematical results in the manuscript. Special thanks to Thomas F. Hansen for his guiding hypotheses on these questions. Funding was provided by the John Templeton Foundation (Grant ID: 61866).

