## Supplementary Appendix for "Universal genomic constraints in the evolvability of thermal physiology"

### Simulation Sensitivity Experiments

#### Methods

We tested the sensitivity of modifying several simulation parameters on rates of evolution. First, the base model $T_b$-genetic (with five loci affecting $T_b$) with no $T_{env}$ was held constant as a reference, and we modified the following parameters individually in separate simulations: either mutation rate or mutational effect size was increased by a factor of 10, or the breadth of the protein thermal performance curve was narrowed by multiplying the Q10 values by 10 (resulting in a relative fitness of 0.94 if $T_b$ was 1°C higher than the $T_{opt}$ of only a single protein). We performed simulations across the same numbers of proteins as in our base simulations, with the exception of experiments increasing the mutation rate, in which we did not analyze models with 100 proteins due to computation limitations. Fold change in rates for sensitivity simulations were interpreted relative to rates in the reference model.

We then tested the sensitivity of our model including a 1°C shift in $T_{env}$ to the same modifications to mutation rate, effect size, and protein performance curve width. The $T_b$-genetic model (with five loci affecting $T_b$) and strong costs of thermoregulation on $T_b$ (standard deviation of 1.0) was held constant as a reference. For experimental simulations in which individual parameters were modified, we recorded the variance in mean $T_b$ among replicates and the $t_{1/2}$, comparing the fold-change with respect to the reference model.

#### Results

In sensitivity experiments without any $T_{env}$, increasing the mutation rate by a factor of 10 in our $T_b$-genetic model had a relatively stable effect on the rate of evolution (i.e., variance across replicates) across greater numbers of proteins, increasing the rate 6.0–7.7 fold (Fig. S1A). Similarly, narrowing the width of the TPC (multiplying Q10 by 10) had a stable effect across numbers of proteins, decreasing the evolutionary rate by 3.2–5.0-fold (Fig. S1A). Increasing the effect size of mutations by 10x, on the other hand, had diminishing effects on the rate of evolution as the number of proteins increased, accelerating the rate of evolution 14.9-fold for a single protein but only 3.7-fold for 100 proteins (Fig. S1A).

When a 1°C shift in $T_{env}$ was included, the variance in final $T_b$ across replicates in sensitivity simulations were similar to the baseline model with most experimental modifications (Fig. S1B). The direction of the effect of modifications depended on the number of proteins, but experiments resulted in only 0.9–2.0-fold change in variance for any modification or number of proteins. However, half-lives were reduced across all sensitivity simulations, suggesting each of the modifications increased the evolvability of $T_b$ given up to 100 proteins (Fig. S1C). In many cases, typically with fewer proteins, half-lives could not be calculated because $T_b$ was shifted completely to the new optimum and slightly exceeded the new $T_{env}$. The magnitude of the effect varied across the numbers of proteins and parameters modified. For the mutation rate and effect sizes, half-lives tended to decrease with greater numbers of proteins, as more mutational input or stronger effect sizes allowed proteins to more effectively respond to directional selection. Narrowing the protein performance curve, however, showed diminishing effects on half-lives at higher numbers of proteins. With narrower performance curves, the initial population response to the shift in $T_{env}$ was dramatically decreased, and the decoupling of the evolution of $T_b$ and proteins was constrained, particularly with higher numbers of proteins (Fig. S3).

#### Correlated Progression model as stochastic differential equations

##### Multivariate Ornstein-Uhlenbeck approximation of the $T_b$-genetic model

We compare our simulation results to analytical solutions derived from analytical approximations of the correlated progression model. Specifically, we approximate the conditions of the correlated progression model under two scenarios 1) the $T_b$-genetic model and 2) the $T_b$-emergent model:

First, we estimate the expectation of correlated progression under a multivariate Ornstein-Uhlenbeck (OU) model for the $T_b$-genetic model. Unlike our simulations, the multivariate OU model assumes Gaussian fitness functions for the costs of thermoregulation and individual protein performance. Under this model, we describe the evolution of a set of coupled stochastic differential equations tracking the body temperature of the environment, $T_b$, and the individual thermal optima of $K$ individual proteins, with individual protein optima indexed by i, $T_{opt,i}$:

$$dT_b = -\alpha_0(T_b(t) - \theta_{T_b})dt - \sum \alpha_i(T_b(t) - T_{\text{opt}_i}(t))dt + \sigma_0 dW_0 \quad (1)$$

$$dT_{\text{opt}_i} = -\alpha_i(T_{\text{opt}_i} - T_b(t))dt + \sigma_i dW_i \quad (2)$$

These equations explain our two-phase evolution evident in Figure 3 of the main text and the eventual slowdown present in adaptation to a new optimum under the $T_b$-genetic model via correlated progression. Specifically, body temperature change evolves as a result of three terms of the stochastic differential equation (SDE; Eq. 1). First, the costs of thermoregulation pull body temperature toward its natural selection optimum, $\theta_{T_b}$. The second term describes the cumulative pull on body temperature toward values that align with individual protein optima. Finally, the third term describes a stochastic Wiener process. The second coupled equation (Eq. 2) describes the evolution of an individual protein's optimum pulled toward the body temperature obtained through thermoregulation. This captures a key feature of correlated progression, where individual proteins are weakly pulled toward body temperature, but body temperature is pulled as the cumulative sum toward the average thermal optimum of each protein. If we consider the deterministic dynamics of this process from a starting point where all $T_{opt}$ and $T_b$ are set at their ancestral optimum, followed by a displacement to a new natural selection optimum, we recover our two-phased adaptive process apparent in simulations (Fig. 4, main text). First, $T_b$ displaces to balance the costs of thermoregulation against the cumulative costs of maladapted proteins—quickly reaching an equilibrium point. However, the population now experiences fitness load resulting from $T_b$ neither being on its natural selection optimum, nor being centered on the thermal optimum of the average protein. Increasing fitness further requires evolution of individual protein optima, which are only weakly pulled toward body temperature. As more proteins are included in the fitness function, the difference between how much $T_b$ is restrained (2nd term of Eq. 1) and how much each individual protein is pulled toward body temperature (Eq. 2) is directly proportional to the number of proteins. This asymmetry means that selection on individual protein optima will be quite weak and evolutionary change will exponentially slow with number of proteins (Fig. 3, main text).

In adaptive processes, we expect that the rate of adaptation will be directly proportional to the additive genetic variance and inversely proportional to the width of the fitness function. Furthermore, we simplify to consider only the deterministic dynamics of adaptation. Under deterministic processes ($\sigma_0^2 = \sigma_i^2 = 0$), all proteins will respond the same to adaptive conditions, and we can simplify the multivariate OU process to a bivariate process:

$$dT_b = -\alpha_0(T_b(t) - \theta_{T_b})dt - K\alpha_i(T_b(t) - T_{\text{prot}}(t))dt \tag{3}$$

$$dT_{\text{prot}} = -\alpha_i(T_{\text{prot}} - T_b(t))dt \tag{4}$$

We consider this model where the rate of adaptation, $\alpha$, is proportional to a traits additive genetic variance, $G$, divided by the width of the Gaussian fitness function $\omega^2$. This gives:

$$dT_b = -\frac{G_{T_b}}{\omega_{T_b}^2}(T_b(t) - \theta_{T_b})dt - K\frac{G_{T_b}}{\omega_{T_{prot}}^2}(T_b(t) - T_{\text{prot}}(t))dt \tag{5}$$

$$dT_{\text{prot}} = -\frac{G_{T_{prot}}}{\omega^2_{T_{prot}}}(T_{\text{prot}} - T_b(t))dt \quad (6)$$

Where $G_{T_b}$ is the additive genetic variance of body temperature, $\omega^2_{T_b}$ is the width of the Gaussian fitness function imposed by the environment (i.e. the costs of thermoregulation), $G_{T_{prot}}$ is the additive genetic variance of the proteome optimal temperature (the average $T_{opt}$ across proteins), and $\omega^2_{T_{prot}}$ is the width of the Gaussian thermal performance curve of each individual protein around its optimum (which is assumed to be equal across proteins).

We use simulations from SLiM with a Gaussian thermal performance curve to validate these results using numerical simulations of a bivariate Ornstein-Uhlenbeck process. Because OU models assume symmetrical fitness functions for proteins, whereas our original SLiM simulations used asymmetric performance curves, we reran $T_b$-genetic simulations using Gaussian protein fitness functions with a SD of 16.5 to approximate the width and curvature of our original thermal performance curves. These simulations included five loci coding for body temperature and our "weaker" organismal costs of thermoregulation (SD = 2.0). We obtain $G_{T_b}$ from simulated runs as the mean additive genetic variance of body temperature and $G_{T_{prot}}$ as the mean additive genetic variance over time and across proteins. That is, to obtain starting values for the the rate of adaptation, $\alpha$, final phenotypic variance across individuals in each simulation was divided by the squared SD of the fitness function ($2.0^2$ and $16.5^2$ for $T_b$ and $T_{prot}$, respectively). We calculated $G_{T_b}$ and $G_{T_{prot}}$ for each simulation and fit separate OU models to estimate the range of expected values. Across simulations, these approximations give good fit to the deterministic dynamics observed in our simulations (Fig. 6, main text) across proteome sizes.

#### Mutational target size in the $T_b$-emergent model

The constraints on thermal evolution were further examined by considering the $T_b$-emergent model, where body temperature is equated to the average value of $T_{opt}$ across all proteins—essentially allowing an organism to assess the performance of the proteome and optimize its physiological body temperature to match. Under this architecture, we consider a population with a perfectly "coordinated" proteome and body temperature where all $T_{opt,1} = ... = T_{opt,K} = T_b$ for all $i$. We then consider whether a new mutation at a given locus would be selectively favored under the $T_b$-emergent model when the environmental optimum for body temperature is shifted such that $T_{env} \neq \theta_{T_b}$. We define these conditions where $\Delta f_{env}$ is the change in fitness due to the costs of thermoregulation (proportional to $|T_b - \theta_{T_b}|$), and $\Delta f_{prot}$ is the change in fitness due to the cumulative effects on fitness of displacing $T_b$ from individual protein optima. A mutation that allows a change in $T_b$ toward the adaptive optimum $\theta_{T_b}$ is then favored if:

$$\Delta f_{env} > -\Delta f_{prot} \tag{7}$$

A new mutation of effect size $\delta$ in protein $i$ that shifts its optimum from $\Delta T_{opt,1} = \delta$ will shift body temperature equal to:

$$T_b^* = \frac{\delta + \sum_1^K T_{opt,i}}{K} = \frac{\delta}{K} + T_{b,0} \tag{8}$$

We ask when the mutant phenotype $T_b^*$ is favored for a given value of $K$. $\Delta f_{env}$ and $\Delta f_{prot}$ were considered in our simulations to follow organismal and the product of independent protein thermal performance curves, respectively. If $T_b < T_{opt}$, these can alternatively be approximated as a linear functions, or considered to follow Gaussian functions as we illustrated for the multivariate OU model. For $f_{prot}$, we make a simplifying assumption that individual protein's TPCs are Gaussian, that all $T_{opt,i} = T_{b,0}$ for $i = 1$ to $i = K - 1$, and that $T_{opt,K} = T_{b,0} + \delta$. This means the fitness of the mutant is:

$$f_p = (Q_K e^{-(\frac{\delta}{K} - \delta)^2/(2\omega_p^2)} \times \prod_{i=1}^{K-1} Q_i e^{-(\delta/K)^2/(2\omega_p^2)}) \tag{9}$$

$$= \prod_{i=1}^{K} Q_i e^{-\frac{\delta^2}{2\omega_p^2}(1-\frac{1}{K})} \tag{10}$$

Where $Q_i$ is a normalizing constant. Standardizing to relative fitness where fitness is maximized when all proteins are coordinated with $T_{opt,i} = T_b$, drops this term from the equation. The change in $f_p$ of the mutant evolving from from $T_{b,0}$ to $T_{b,0} + \frac{\delta}{N}$ is thus:

$$\Delta f_p = e^{-\frac{\delta^2}{2\omega_{prot}^2}(1-\frac{1}{K})} - 1 \tag{11}$$

When $T_b < T_{th}$, $\Delta f_{env}$ for our environmental thermal performance curves is equal to:

$$\Delta f_{env} = C(e^{\nu(T_{b,0}+\delta/K)} - e^{\nu T_{b,0}}) = Ce^{\nu T_{b,0}}(e^{\delta/K} - 1) = \beta(e^{\delta/K} - 1) \tag{12}$$

where $\nu = log(Q_{10})/10$ and $C$ is a normalizing constant. Alternatively, we consider a linear approximation of a TPC when $T_b < T_{opt}$, where $\Delta f_{env}$ is then:

$$\Delta f_{env} = \beta\frac{\delta}{K} \tag{13}$$

Where $\beta$ is the slope of the increase in thermal performance approaching the maximum value. The conditions for the spread of the mutant allele from our initial conditions are therefore:

$$\beta(e^{\delta/K} - 1) > e^{-\frac{\delta^2}{2\omega_{prot}^2}(1-\frac{1}{K})} - 1 \tag{14}$$

or for a linear approximation:

$$\beta\frac{\delta}{K} > e^{-\frac{\delta^2}{2\omega_{prot}^2}(1-\frac{1}{K})} - 1 \tag{15}$$

In Fig. S7B and S7C we plot the relationship between $\Delta f_{env}$ and $\Delta f_p$ and mutant effect size, $\delta$ for various values of $K$. This demonstrates that increasing proteome size has two major effects. First, it decreases mutational target size. This is because mutations that increase an individual protein's thermal optimum toward the natural selection optimum set by the environment are overwhelmed and offset by the cumulative effects of uncoordinated proteins around the new body temperature. This explains why our simulations that increased mutational effect size did not increase the rate of evolution with increasing proteome size (Fig. 5a, main text), as it results in the vast majority of new mutations being selectively disfavored. Second, as the mutational target size decreases with increasing proteome size, the strength of natural selection on those new mutations likewise decreases (Fig. S7D). This means these favorable mutations approach neutrality as the target size decreases, lengthening their expected time to fixation and the waiting time of a new substitution. These two effects reveal the overwhelming effects of sign epistasis and correlated progression on the adaptive process for body temperature. Solving for the intersection point between our two functions $\Delta f_{env}$ and $\Delta f_{prot}$ reveals an asymptotically log-linear relationship, allowing extrapolation to proteomic scales (Fig. S7C):

$$\Delta f_{env} = -\Delta f_{prot} \tag{16}$$

$$\beta \frac{\delta}{K} = 1 - e^{-\frac{\delta^2}{2\omega_{prot}^2}(1-\frac{1}{K})} \tag{17}$$

Taking the log of both sides

$$\log \beta + \log \delta - \log K = \log \left( 1 - e^{-\frac{\delta^2}{2\omega_{prot}^2}(1-\frac{1}{K})} \right) \tag{18}$$

For the right side of the equation, when $K \to \infty$, the term $1 - 1/K$ approaches 1. Setting $a = 1/(2\omega_{prot}^2)$ and a Taylor series expansion for $\log(1 - e^{-a\delta^2})$ results in:

$$\log \left(1 - e^{-a\delta^2}\right) \approx \log(a\delta^2 - \frac{1}{2}a^2\delta^4) = \log(a\delta^2) + log(1 - \frac{1}{2}a\delta^2)) \tag{19}$$

Using the approximation $\log(1+\epsilon) \approx \epsilon$ for small $\epsilon = -\frac{1}{2}a\delta^2$ results in:

$$\log(a\delta^2) + log(1 - \frac{1}{2}a\delta^2) = \log a + 2\log\delta - \frac{1}{2}a\delta^2 \tag{20}$$

Substitution back into equation 18 results and rearranging results in:

$$\log\beta + \log\delta - \log K \approx \log a + 2\log\delta - \frac{1}{2}a\delta^2 \qquad (21)$$

$$\log\delta \approx (\log\beta - \log a) - \log K + \frac{1}{2}a\delta^2 \qquad (22)$$

At small $\delta$, $\frac{1}{2}a\delta^2 \to 0$, and setting $D = \log\beta - \log a$ reveals a simple log-linear relationship between $\log\delta$ and $\log K$ with a slope of -1 as $\delta \to 0$ for large $K$:

$$\log\delta \approx -\log K + D \qquad (23)$$

In other words, the maximum mutational effect size that is selectively favorable scales inversely with $\frac{1}{K}$, such that for proteomic scales (e.g. K = 10,000 - 30,000), we expect the mutational target size that is favored would likely be of the order $10^{-3}$ to $10^{-4}$ $°C$ with selection differentials well below the drift-fixation threshold of 1/(2*Ne)((Fig. S7D)), resulting in fixation rates and probabilities similar to that of neutral drift.

Using these results, we can approximate the long-term dynamics of evolution under the $T_b$-emergent model. Here, we assume that the mutational target size derived under these results remains constant when there is standing genetic variation in protein optima across the proteome. In other words, that standing genetic variation only imposes a fixed genetic load under equilibrium without dramatically altering mutational target size. With this assumption, we explore the consequences of reduced mutational target size on adaptive and non-adaptive evolutionary dynamics. Under our model of adaptive evolution, only nearly neutral mutations can go to fixation, with probabilities and times to fixation equal to that of genetic drift when $K$ is large. However, the mean of this distribution is shifted slightly toward the adaptive optimum. This means that the process will behave as Brownian Motion with a trend, with the trend parameter equal to the mean of the distribution of nearly-neutral mutations, $\mu_{nn}$, and the variance of that distribution being represented as $\sigma^2_{nn}$. Under purely neutral processes, the rate of new substitutions in the population is $\nu K$ where $\nu$ is the per locus mutation rate (1). The effective population size and ploidy, $2N_e$, cancels out of the equation because it appears in both the numerator ($2N_e$ gene copies in which potential mutations can occur) and the denominator (the probability of fixation of a new mutation is $\frac{1}{2N_e}$). However, not all substitutions will be neutral—some will be strongly deleterious. Only a fraction of those mutations will be close enough $\mu_{nn}$ to be nearly-neutral, and the proportion of these nearly-neutral mutations as a fraction of the total mutation rate is represented by $p_{nn}$. Thus, for our scenario of an adaptive shift of 1°$C$, we can calculate the amount of time ($t$) to get halfway (0.5°C) as:

$$t(0.5°C) = g \times \frac{0.5}{K\nu\mu_{nn}p_{nn}} \tag{24}$$

Where $g$ is the generation time. Additionally, without selection, $\mu_{nn} = 0$ and the process evolves as an unbiased random walk with step variance equal to:

$$\sigma^2_{bm} = g \times K\nu\sigma^2_{nn}p_{nn} \tag{25}$$

What is the proportion of nonsynonymous substitutions that fall within the neurly-neutral mutational window? (2) compiled a database of the thermodynamic properties of 1,626 single-nucleotide polymophisms in about 93 small proteins and measured the change in melting temperature and free energy as measures of thermodynamic stability. Of those 1,626 proteins, 58 had no detectable effect on melting temperature ($T_m$) or free energy, whereas most had relatively large effects ($|\Delta\bar{T}_m| = 4.2$°C, 95% CI: 0, 15.74 °C). Here, we consider the estimate of $p_{nn} = 58/1,626 = 0.036$. as a likely overestimate of the proportion of nearly-neutral mutations, as the limits of detection of change in $T_m$ and free energy likely exceed the limits of our nearly-neutral window $s << \frac{1}{2N_e}$ for biologically realistic values of $N_e$.

Using numerical Monte Carlo simulations, we simulate a range of parameters under our scenario outlined in Fig. S7. We consider $\log_{10} N_e \sim logNorm(5, 0.15)$, $\log_{10} \nu \sim U(-8, -5)$, $g \sim logNorm(log(5yrs), 0.25)$, $K \sim U(10000, 30000)$, and $p_{nn} \sim U(0, 0.036)$ to obtain a distribution of plausible rates of adaptation and genetic drift (Fig S8). We then compared these to macroevolutionary estimates of these parameters from (3) estimated for preferred body temperature from the adaptation-inertia model of (4) in mammals, birds, and squamates (Fig. 3B in main text). While the uncertainty in parameters generates a large region of potential values (especially when considering the logarithmic scale used), we nevertheless recover an average value that is remarkably close to empirical estimates, and well within the region of plausible values (Fig. 3B in main text). This suggests that regardless of the specific conditions, it is at least plausible that the correlated progression model and its effect via sign epistasis on decreasing the mutational target size to nearly-neutral mutational effects on body temperature is more than capable of explaining both neutral and adaptive dynamics of thermal evolution observed at macroevolutionary scales.

#### Supplementary Figures

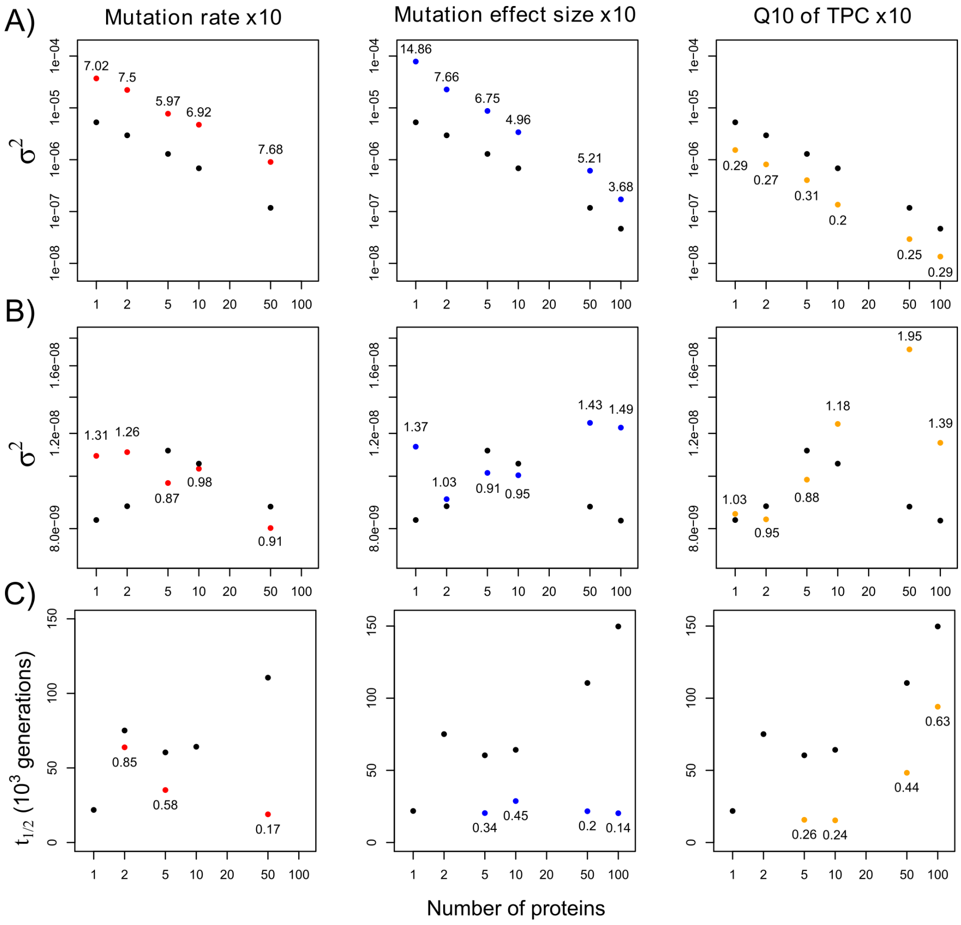


Figure S1: Experimental sensitivity simulations for the effects of increasing the mutation rate, mutational effect sizes, or decreasing the Q10 (i.e., width) of the thermal performance curve (TPC) by factors of 10. Simulations of the baseline models (black circles) were the $T_b$-genetic model with five loci for Tb, and either did not include any environmental temperature (A), or they included a 1°C shift in environmental temperature partway into the simulation (B – C). A) Values represent variance ($\sigma^2$) in final mean population $T_b$ across 100 simulated replicates. Colored circles are experimental values, and the associated number is the fold change relative to the base model. Panels (B – C) are the variance and half-lives, respectively, for the model including a 1°C shift in environmental temperature. Missing experimental values for half-lives resulted if the average final $T_b$ exceeded the new environmental temperature.

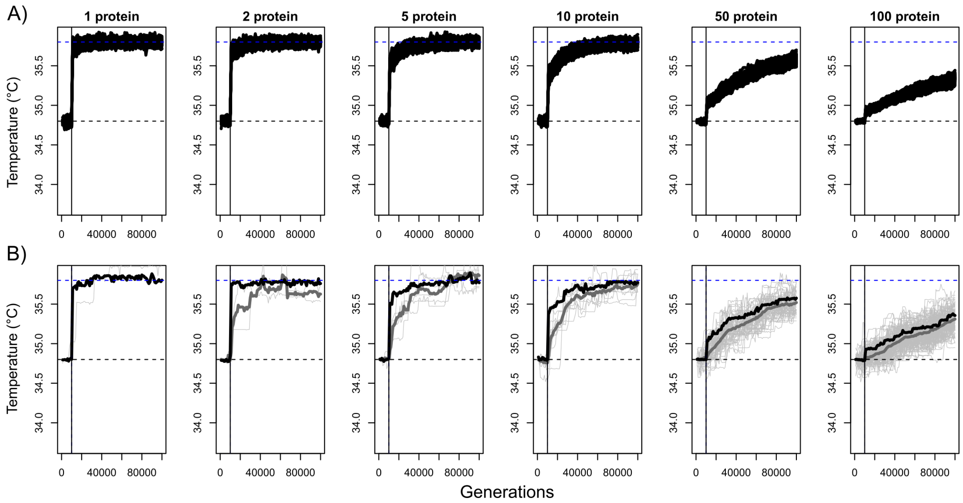


Figure S2: Experimental simulations with a 1°C increase in $T_{env}$ and the Q10 of the protein thermal performance curves (i.e. the width parameter) decreased by a factor of 10x, causing stronger selection from individual proteins. Simulations used the $T_b$-genetic model (five loci for $T_b$) with our strongest cost of thermoregulation pulling $T_b$ towards the new $T_{env}$. A) Results for average population $T_b$ for 100 simulated replicates (black lines). B) An example simulation, also showing protein optima (light gray) and the average optimum across proteins (thick gray).

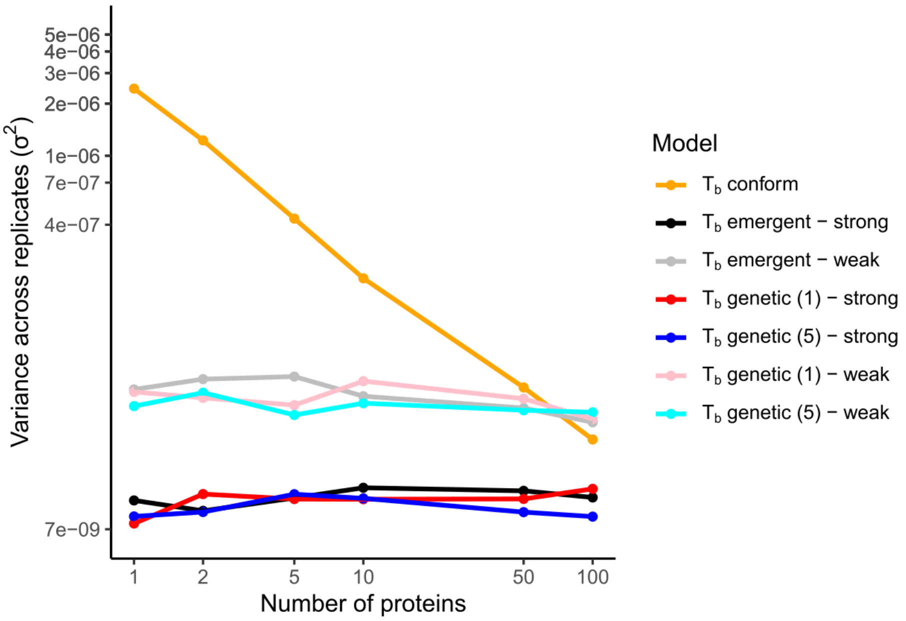


Figure S3: Line plot of final $T_b$ variation across replicates for simulations including a 1°C shift in environmental temperature. These results differed little from those with a static environmental temperature (Fig. 2B in main text). Note the log10-transformed x- and y-axes shown with original units. Legend values in parentheses for $\mathrm{T_b}$-genetic models indicate whether one or five loci were used to control organismal $T_b$. "Strong" and "weak" refer to the cost of thermoregulation (i.e. whether fitness functions for organisms mismatching environmental temperature used a mean-zero normal distribution with a standard deviation of one or two)

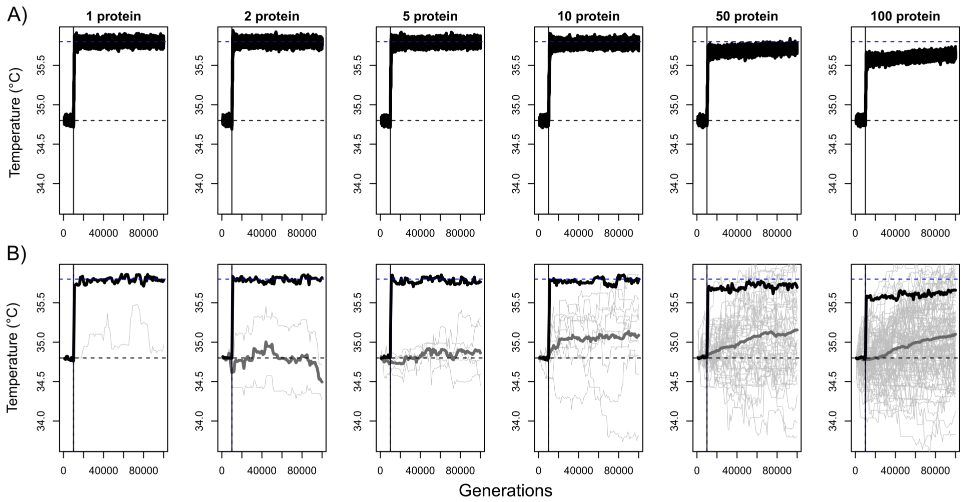


Figure S4: Simulations using the $T_b$-genetic model (five loci for $T_b$) with our strongest cost of thermoregulation pulling $T_b$ towards the new Tenv. A) Results for average population $T_b$ for 100 simulated replicates (black lines). B) An example simulation, also showing protein optima (light gray) and the average optimum across proteins (thick gray).

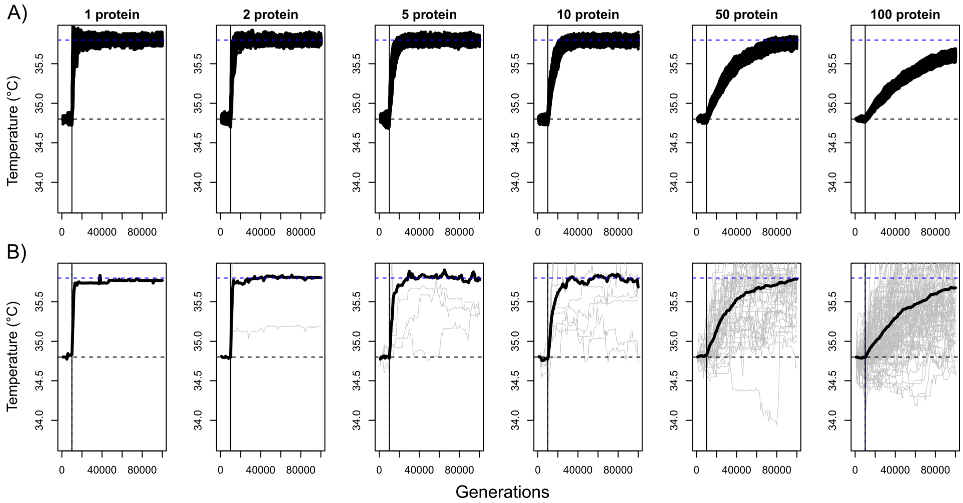


Figure S5: Simulations using the $T_b$-emergent model with our strongest cost of thermoregulation pulling $T_b$ towards the new $T_{env}$. A) Results for average population $T_b$ for 100 simulated replicates (black lines). B) An example simulation, also showing protein optima (light gray).

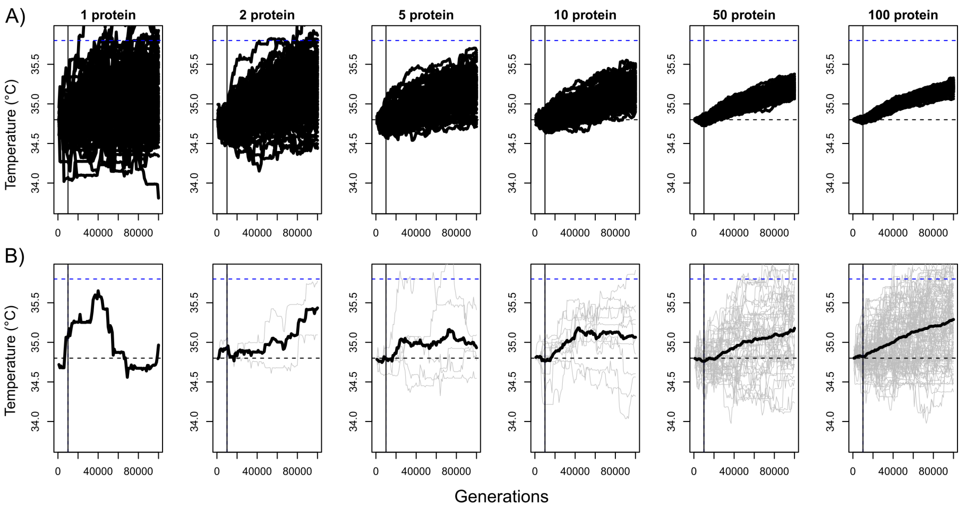


Figure S6: Simulations using the $T_b$-conform model and a 1°C increase in $T_{env}$. A) Results for average population $T_{opt}$ (there is no $T_b$ per se in the $T_b$-conform model; see text) for 100 simulated replicates (black lines). B) An example simulation, also showing protein optima (light gray).

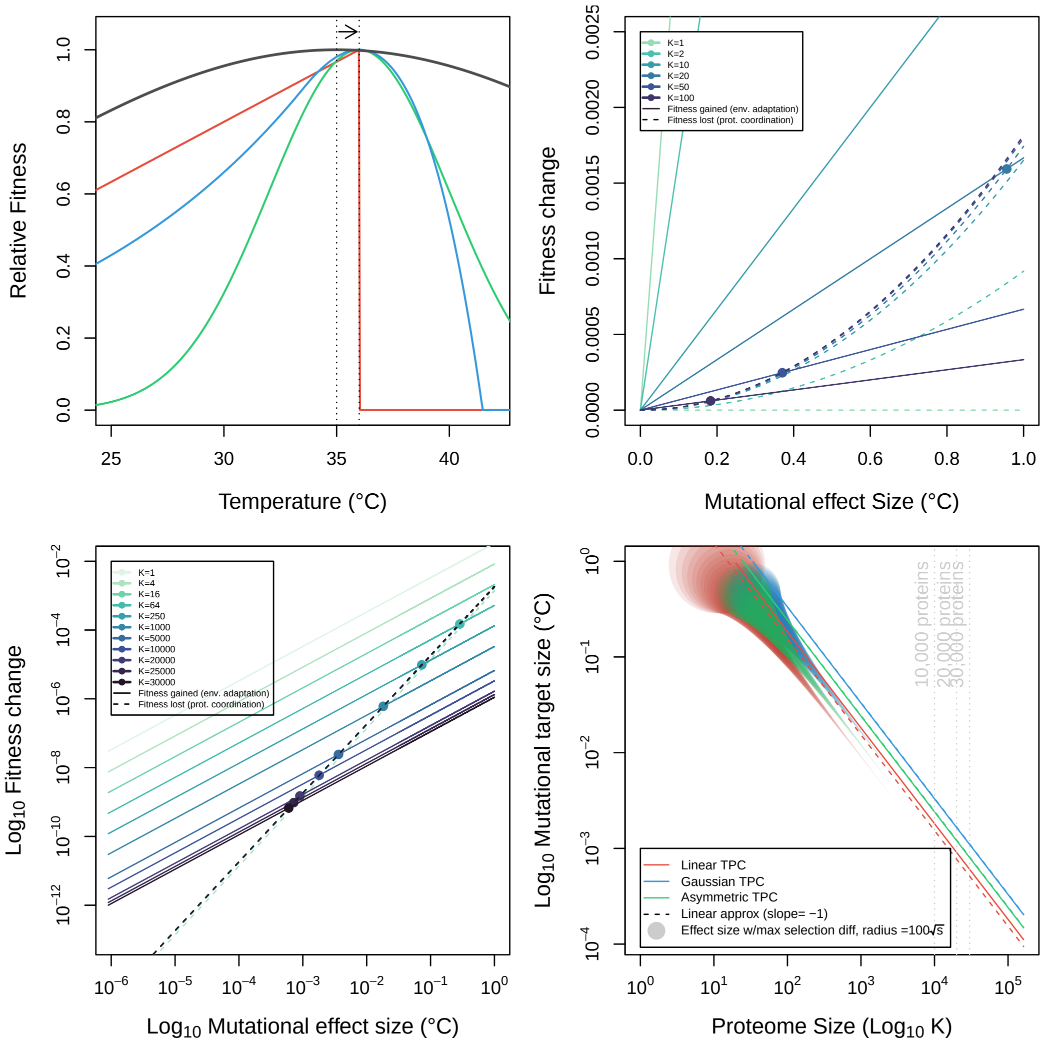


Figure S7: $T_b$-emergent model effects on mutational target size through sign epistasis of effects. A) Thermal performance curves for examined scenarios. In all cases, protein thermal performance curves across all $K$ loci are initiated in a fully-coordinated state with all thermal optima at 35 $°C$, with Gaussian curves and $\sigma^2_{prot} = 16.5^2$ (black). The costs of thermoregulation set by environmental temperature follow asymmetric (blue, $Q_{10}$= 2.356, d=0.016), Gaussian (green, $\omega^2_{env} = 4^2$) or linear (red, $\beta_{env} = 1/30$) curves with thermal optima at $T_{opt} = 36°C$. Curve parameters were chosen to be roughly similar in their overall magnitude over the interval of the considered shift, from $35 \rightarrow 36°C$. B) Fitness gained (solid lines) against fitness lost (dotted) lines with color indicating proteome sizes (darker colors are larger proteomes). When $K = 1$, there are no costs of uncoordinated proteomic thermal optima. As $K$ increases, mutational effect sizes shift $T_b$ by decreasing magnitudes in the $T_b$-emergent model, resulting lower fitness gains while increasing proteomic costs, until the costs intersect with the fitness benefits (solid points). The point of this intersection results in decreasing mutational effect sizes with increasing $K$. C) Same plot as (B) but on the logarithmic scale. As $K$ increases, the costs to the proteome converge to a single line, while the benefits keep decreasing, such that only mutational effects with very small effects (e.g. $10^{-3} - 10^{-4}$ and very small fitness differentials ($s << 10^{-8}$) are selectively favored, placing these mutations in the nearly-neutral drift range. D) Solid lines indicate numerical evaluations of the maximum selectively favored mutational effect size for increasing values of $K$, with the dotted lines representing linear functions with slope -1 and intercepts equal to the fitted values, confirming our analytical results. Line colors correspond to linear (red), Gaussian (green) and asymmetric (blue) costs to thermoregulation as in (A). Points below the solid lines of the same color indicate the value of mutational effect that maximizes the selection differential in favor of $T_b$ adaptation, with the size of the point proportional to $100 \times \sqrt{s}$. Note that the size of the selection differential quickly decreases to near 0 (nearly neutral) as we approach realistic proteomic sizes.
